# Using solid-state NMR to understand the structure of plant cellulose

**DOI:** 10.1101/2024.08.24.609305

**Authors:** Rosalie Cresswell, Parveen Kumar Deralia, Yoshihisa Yoshimi, Tomohiro Kuga, Alberto Echevarría-Poza, W. Trent Franks, Steven P. Brown, Ray Dupree, Paul Dupree

## Abstract

The structure of plant cellulose microfibrils remains elusive, despite the abundance of cellulose and its utility in industry. Using 2D solid-state NMR of ^13^C labelled never-dried plants, six major glucose environments are resolved which are common to the cellulose of softwood, hardwood and grasses. These environments are maintained in isolated holo-cellulose nanofibrils, allowing more detailed microfibril characterisation. We show there are only two glucose environments that reside within the microfibril interior. These have the same NMR ^13^C chemical shifts as tunicate cellulose Iβ centre and origin chains, with no cellulose Iα being detected. The third major glucose site with a carbon 4 chemical shift near 89 ppm, previously assigned to the crystalline microfibril interior, is now shown to be one of four surface glucose environments. The NMR peak widths of all four surface glucose environments are similar to those of the core indicating that their glucose local order is comparable; there is no significant ‘amorphous’ cellulose in the microfibrils. Consequently, the ratio of the carbon 4 peaks at ∼89 and ∼84 ppm, which has often provided a sample cellulose crystallinity index, is not a meaningful measure of crystallinity or the interior to surface ratio. The revised ratio for poplar wood microfibrils is estimated to be 1:2, which is consistent with a cellulose Iβ 18-chain microfibril having 6 core and 12 surface chains, although other microfibril sizes are possible. These advances change substantially both the interpretation of solid-state NMR studies of cellulose and the understanding of cellulose microfibril structure and crystallinity.

## 1. Introduction

Cellulose is the most abundant natural polymer in the world.^1^ It forms a major component in the cell walls of plants providing much of their mechanical strength.^2^ Its properties are important for the pulp and lumber industries and more recently cellulose has become a major contender as a sustainable alternative to fossil fuels for producing materials, films and biofuels.^3–5^ Despite the large industrial interest in cellulose, its structure in native plant cell walls remains elusive.

Cellulose is a β-1-4-linked polymer of D-glucose in long chains of perhaps 10,000 units.^6^ These glucan chains have a 2-fold helical structure which allows them to crystallise via both stacking interactions and a network of hydrogen bonding to form long microfibrils in plant cell walls.^7^ Estimates of the diameter of these cellulose microfibrils are between 3-5 nm.^8^ In recent work, based on biophysical measurements and advances in understanding the biosynthesis, it is believed that a microfibril is formed from 18-24 glucan chains.^9–15^ The arrangement of these chains in the microfibril into several stacked sheets also remains unknown, but computational studies have suggested some possible habits.^8,16–18^

Cellulose Iα and Iβ are well known forms of crystalline cellulose and have been characterised using both diffraction techniques and solid-state NMR.^19–23^ Cellulose Iα has a P1 triclinic structure and has non-equivalent glucose residues in a cellobiose unit within the same chain, whereas cellulose Iβ has a P2_1_ monoclinic structure and its two non-equivalent glucose residues are in alternating sheets of glucan chains, called centre and origin.^19,20^ It is known that cellulose from bacteria and some algae have predominantly a cellulose Iα structure, whilst tunicate cellulose, which consists of large crystals, is cellulose Iβ. Although it is possible to obtain high resolution X-ray and neutron diffraction patterns for these larger crystalline forms of cellulose, this is not achievable on native plant cellulose.^24–26^ The very thin plant microfibrils result in low resolution diffraction patterns such that atomic resolution cannot be achieved.^26–28^

Solid-state magic angle spinning (MAS) NMR has been used for studying cellulose since the 1980s, mainly using 1D ^13^C cross-polarisation (CP) experiments.^29–31^ It is particularly useful as, in contrast to diffraction, it does not require long-range order, and samples can be analysed in situ. Nevertheless, the thin nature of the plant microfibrils means that the surface glucose residues contribute to the NMR spectra far more than in large cellulose crystals of tunicates or fibrils of bacteria, partially obscuring the signals of the microfibril core and complicating attempts to fully assign the spectra. Furthermore, spectra of whole plant samples contain non-cellulosic spectral peaks, as well as some broader components potentially arising from a so-called amorphous cellulose component, that fall in the same region as those of microfibrillar cellulose.^32,33^ The cellulose microfibrils also interact with other components in the cell wall which is likely to influence the ^13^C chemical shifts of surface glucose residues.^34–36^ Within these limitations, it is widely thought, based on 1D NMR assignments and the fitting of X-ray diffraction patterns, that cellulose of higher plants is a mixture of mainly cellulose Iβ and some cellulose Iα.^28^ The poor match to the ^13^C NMR chemical shifts of these two cellulose I allomorphs has also led several studies to suggest that plant cellulose could have its own distinct structure.^37,38^

In NMR spectra of cellulose, the carbon chemical shifts of C4 and C6 of the different glucose units in cellulose are split into two main regions which, following Dupree et al.,^35^ are defined as spectral domain 1 glucosyl residues which have C4^1^ and C6^1^ ^13^C chemical shifts of ∼89 ppm and ∼65 ppm respectively, and spectral domain 2 glucosyl residues which have C4^2^ and C6^2^ ^13^C chemical shifts of ∼84 ppm and ∼62 ppm. Glucose residues with ^13^C chemical shifts in spectral domain 1 have been classed, largely by groups studying cellulosic materials such as pulp, to reflect crystalline cellulose as their shifts are close to those of the crystalline cellulose Iα and Iβ^32,39^ whilst spectral domain 2 is then in turn assigned as arising from amorphous cellulose. This leads to the ratio of signal in these two domains frequently being defined as the crystallinity index (CI) of a sample and is used alongside crystallinity measures from XRD despite large differences in their values.^18,40,41^ In other research groups, domain 1 glucose residues are considered to be interior chains of the microfibril^31,32,42^ and domain 2 glucose residues are believed to be surface glucan chains of the cellulose microfibril.^31,32,39^ Using this alternative assignment has led to the ratio of the two domains being used to provide an estimate of the size of the microfibril and has reignited the debate of how many chains form a microfibril.^9,11,15^ When restricted to 1D ^13^C MAS NMR, it is only possible to resolve residues in these two domains using C4 and C6 shifts, as the C5 shifts overlap with those from C2 and C3, and all the C1 shifts of all glucose residues also overlap.^35^ The C4 cellulose region of the 1D ^13^C NMR spectrum tends to be reasonably clear of contributions from hemicelluloses and pectin and so the ratio of the C4 carbons of the two domains (C4^1^:C4^2^) has been widely used to characterise and compare the cellulose in different plants and cellulosic materials.^43,44^ It has been suggested ^45^ that the major source of this change in ^13^C chemical shift between the two domains seen in carbons 4, 5 and 6 arises from a change in the conformation of the hydroxymethyl group of the glucose residue from *tg* in spectral domain 1 to *gt* or *gg* in spectral domain 2, and this has since been supported by several DFT and NMR studies.^16,46,47^

In recent years, the availability of high field NMR spectrometers and developments in ^13^C labelling of plant cell wall materials (and dynamic nuclear polarisation (DNP)) have enabled access to a wealth of 2D MAS NMR experiments that have allowed unparalleled insight into the structure of plant cell walls and other biological polymer matrices.^48–50^ These studies have made significant strides in understanding hemicellulose structure and interactions with cellulose in both primary (PCW) and secondary cell walls (SCW).^51,52^ Solid-state NMR has been used to identify the 2-fold helical structure of xylan in SCW, and the molecular architecture and water-polymer interactions in softwoods.^34,52,53^ With an aim of understanding the structure of the plant cellulose microfibrils, several distinct glucosyl residues, each with differing ^13^C chemical shifts due to their different local environments, were described by Wang et al. who found similar glucose environments in several plants *(Brachypodium*, *Zea mays*, and *Arabidopsis thaliana*).^43,54^ They suggested cellulose microfibril models based on their understanding that the five domain 1 glucose environments arise solely from sites in the interior chains of the microfibril and the two domain 2 environments lie on the surface.^47,55^ Under this assumption, and using quantitative NMR, they found that cellulose had too many interior relative to surface residues for the favoured microfibril size of 18-24 chains. Additionally, some of the ‘interior’ residues were more distant from water than others which is difficult to reconcile with an 18-24 chain fibril. These discrepancies between the fibril models and NMR data led to the suggestion that the cellulose microfibrils bundle producing a cellulose structure with two layers of interior glucose residues with differing water proximity.^43^

In this work, we assign the ^13^C NMR chemical shifts of the main cellulosic glucose environments found in a range of plant secondary cell wall materials and begin to determine their position in the cellulose microfibril. To achieve this, we studied both poplar wood and xylanase-treated holo-cellulose nanofibrils (hCNFs) prepared from poplar wood using minimal processing to maintain the native cellulose structure.^56^ By removing hemicellulose, lignin, and any amorphous cellulose present we are left with a clearer spectrum with improved resolution which provides more certainty in the assignment of ^13^C NMR chemical shifts for each distinct glucose environment. By comparing with the recently corrected ^13^C NMR chemical shift assignments for cellulose I^23,57,58^ and our spectra of tunicate cellulose we conclusively show that plant cellulose is not a mixture of Iα and Iβ, and that the core of the native cellulose microfibril has ^13^C NMR chemical shifts identical to tunicate cellulose Iβ. We also show that categorising the glucose residues by their chemical shifts into spectral domain 1 or domain 2 does not reflect a split into crystalline and amorphous cellulose nor do they exclusively reflect interior and surface cellulose. In particular, we find that one of the domain 1 glucosyl residue environments is part of a surface glucan chain. (To assist the reader, SI Tables 4 and 5 list key terms and acronyms used in this work.) Taken together, these findings identify and resolve several widely held misconceptions and misinterpretations of solid-state NMR data, and they provide a strong basis to build more coherent models for cellulose in plant cell walls.

## 2. Results

### 2.1. Assigning ^13^C NMR chemical shifts for the glucosyl residue environments in cellulose within plant cell walls

Throughout our work^34,53,59,60^ never-dried samples of many different plants have been studied using ^13^C MAS NMR to understand the structure and interactions of cellulose, hemicellulose, lignin and water in native plant cell walls. The secondary cell wall of poplar wood is relatively simple with only 3 main components: cellulose, xylan and lignin. Fig. 1 shows that the 1D cross polarization (CP) MAS NMR spectrum of the neutral carbohydrate region is dominated by the cellulose signal. The carbon 1 (C1) peak is at ∼105 ppm, C2, C3, C5 overlap in the region between 70-76 ppm and the C4 and C6 carbons are split into two main peaks. The cellulose glucose environments in spectral domain 1 show C4^1^ and C6^1^ peaks at ∼89 ppm and ∼65 ppm respectively and the C4^2^ and C6^2^ spectral domain 2 peaks are at ∼84 ppm and ∼62 ppm respectively. Both these C4 and C6 regions in the 1D CP MAS spectrum are comprised of several peaks since multiple glucose environments contribute to these spectral domains. The C4^2^ region especially is clearly split into two peaks at ∼83.5 and ∼84.5 ppm.

**Figure 1:**
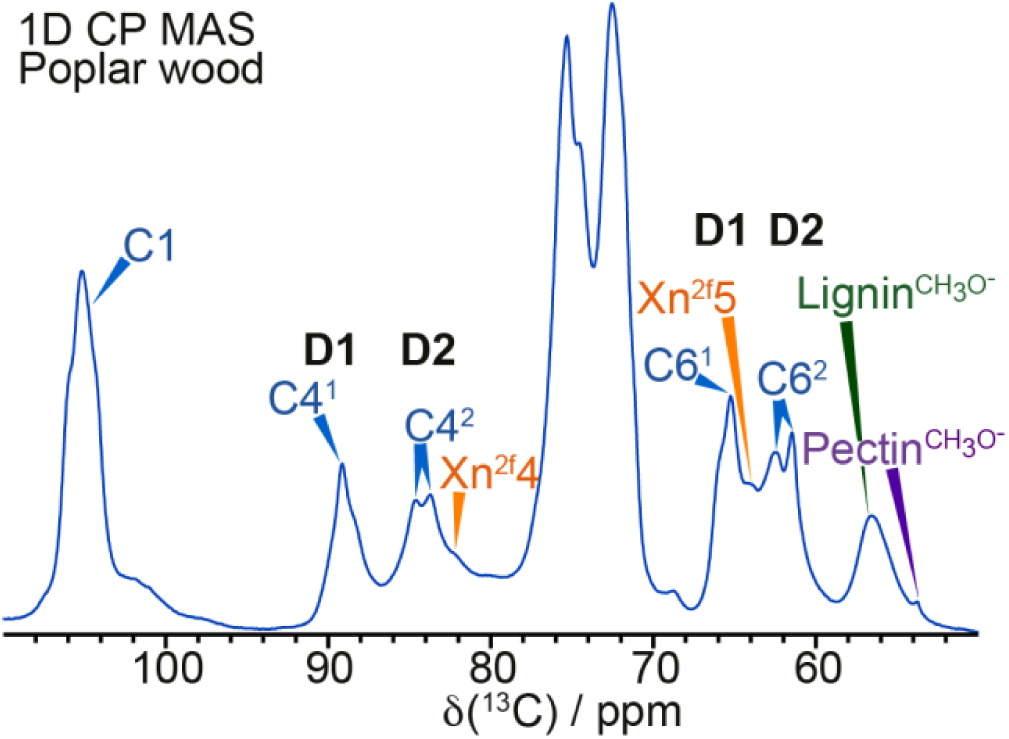
The neutral carbohydrate region of a ^13^C 1D CP MAS NMR spectrum of poplar wood. The spectrum shows the assignments of cellulose (blue), xylan (orange), lignin (green) and pectin (purple). The glucose C4 and C6 peaks in cellulose are split into two regions named spectral domain 1 (D1) and spectral domain 2 (D2). The spectrum was recorded at a ^13^C Larmor frequency of 251.4 MHz and a MAS frequency of 12.5 kHz.

Fully ^13^C labelled never-dried poplar wood enables the structure of plant cellulose to be explored using a wealth of 2D MAS NMR experiments.^34,43,61,62^ The CP MAS refocussed INADEQUATE NMR experiment^63,64^ is a double quantum experiment that correlates two covalently bonded carbons with directly bonded carbons appearing at the same Double Quantum (DQ) shift given by the sum of the respective Single Quantum (SQ) shift of the bonded carbons. Fig. 2 shows the neutral carbohydrate region of a ^13^C CP MAS refocussed INADEQUATE NMR spectrum of poplar wood. The high resolution means that the carbons within each glucosyl residue can be followed through in the 2D spectrum.

**Figure 2:**
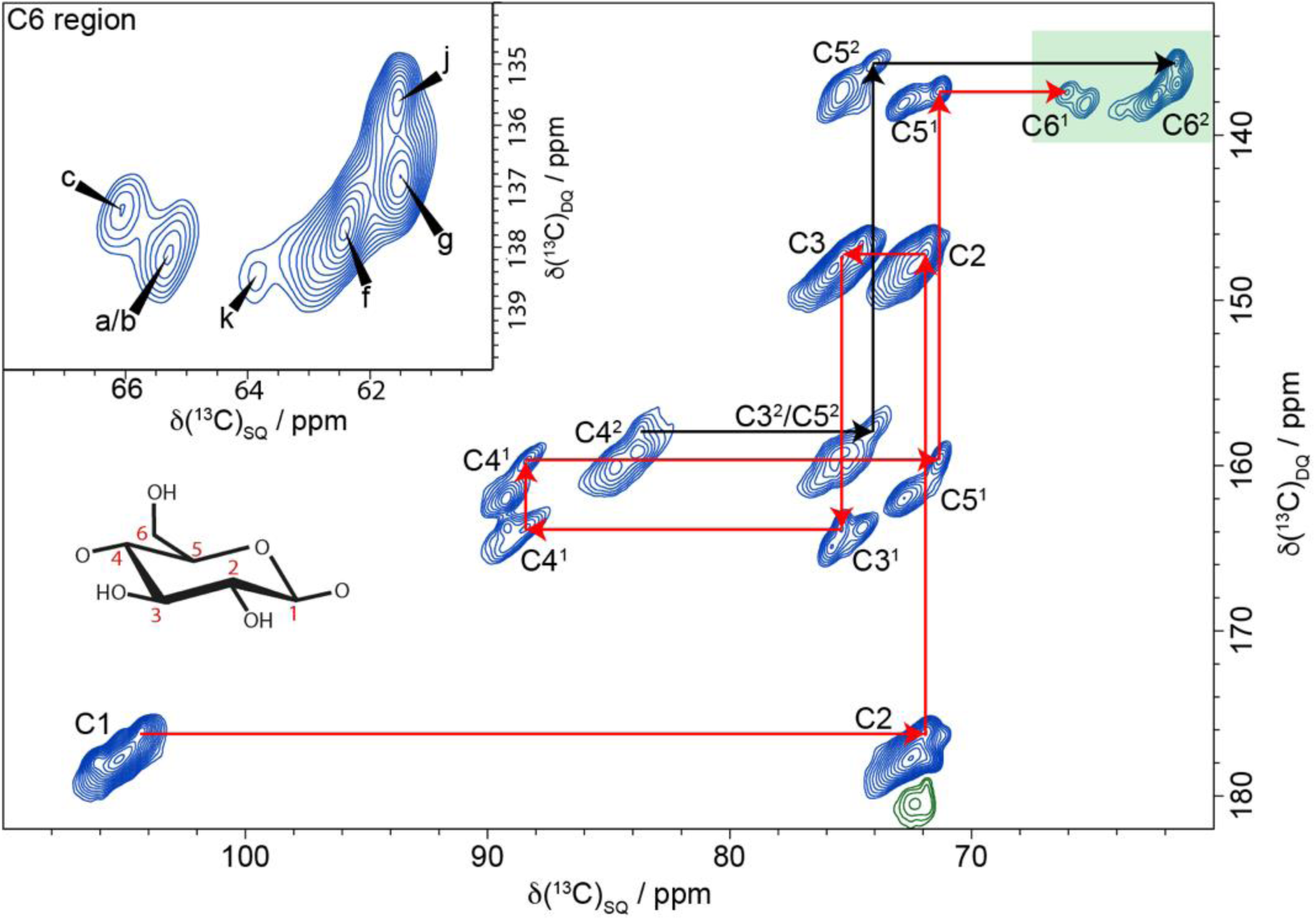
A ^13^C 2D CP refocussed INADEQUATE MAS NMR spectrum of never-dried poplar wood showing multiple glucose environments in both spectral domains 1 and 2. The path of ^13^C carbon chemical shifts for cellulose glucose residue environments c (red) and j (black) are shown as example assignments. **Inset (top left):** The C6 region with the six major glucose environments identified in poplar wood (a,b,c,f,g,j). The spectrum was recorded as in Fig. 1.

This is demonstrated in red for a glucose residue in domain 1 named environment ‘c’, and in black for the C4-C6 region for a residue in domain 2 environment ‘j’. Our naming convention and assignments are based on those of Wang et al.^54^ for cellulose in primary cell walls since many of the cellulose environments we observe have similar values. However, glucose environment j, which is clearly visible for C4, C5 and C6 in the ^13^C CP MAS refocussed INADEQUATE NMR spectrum, is newly reported here. The C6 region of the spectrum (inset in Fig. 2) shows the main glucose environments identified within spectral domain 1 and spectral domain 2. This region also shows a minor domain 2 environment, named k. We found, in contrast to previous work on plant primary cell walls,^54^ that there are just six major glucose environments in this never-dried poplar sample: three (a, b, c) in spectral domain 1 and three (f, g, j) in spectral domain 2.

To make a more complete assignment of the ^13^C NMR chemical shifts of the glucose environments within cellulose, a 30 ms ^13^C CP MAS proton-driven spin-diffusion (PDSD) NMR spectrum was analysed alongside the ^13^C CP MAS refocussed INADEQUATE NMR spectrum. The ^13^C CP MAS PDSD NMR experiment is a through-space correlation experiment for which, with a mixing time of 30 ms, the observed cross peaks are found to only correlate carbons that are within the same glucose ring. The 30 ms CP MAS PDSD NMR spectrum shown in SI Fig. 1 highlights the key regions (C6-C1, C6-C4, C6-C5 and C4-C1) which are particularly useful for identifying different glucose environments. Two further minor glucose environments in domain 1 labelled d and e (which are very minor in our poplar wood sample) have previously been assigned by Wang et al.^54^ Table 1 lists the ^13^C NMR chemical shifts for the six major and the minor glucose environments identified in cellulose of poplar wood.

**Table 1.** NMR chemical shifts of all glucose environments assigned in cellulose of poplar wood.

| Spectral Domain 1 |  |  |  |  |  |  |
| --- | --- | --- | --- | --- | --- | --- |
| Glucose Environment | C1<sup>1</sup> | C2<sup>1</sup> | C3<sup>1</sup> | C4<sup>1</sup> | C5<sup>1</sup> | C6<sup>1</sup> |
| a* (Cellulose Iβ origin) | 106.1<sup>§</sup> | 71.8 | 74.5 | 89.2 | 72.8 | 65.2 |
| b | 105.3 | 72.6 | 75.5 | 89.3 | 72.7 | 65.4 |
| c (Cellulose Iβ centre) | 104.3 | 71.8 | 75.4 | 88.4 | 71.3 | 66.0 |
| *d* | *105.2* | *72.5* | *74.9* | *87.1* | *72.5* | *64.7* |
| *e* | *105.0* | - | *74.7* | *89.8* | *71.1* | *65.3* |
| Spectral Domain 2 |  |  |  |  |  |  |
| Glucose Environment | C1<sup>2</sup> | C2<sup>2</sup> | C3<sup>2</sup> | C4<sup>2</sup> | C5<sup>2</sup> | C6<sup>2</sup> |
| f | 105.2 | 72.4 | 74.4 | 84.5 | 75.3 | 62.4 |
| g | 105.0 | 72.4 | 75.3 | 83.5 | 75.2 | 61.5 |
| j | 105.1 | - | - | 83.3 | 73.9 | 61.4 |
| *k* | - | - | - | *83.8* | *74.3* | *63.7* |

*N.B. The naming convention and assignments are based on those of Wang et al.<sup>54</sup> with additional environments such as j assigned in this work. The minor glucose environments d, e and k are italicised.*
*<sup>§</sup>All <sup>13</sup>C NMR chemical shifts are in ppm with an error of ± 0.1 ppm.*

### 2.2. Glucose environments in cellulose in cell walls of different plants

Having identified the six glucose environments in cellulose of poplar wood, we were interested in the presence of these in the cellulose of other plant species. Over the years we have used high resolution 2D MAS NMR experiments to study a wide range of different plants including monocots, eudicots, and gymnosperms.^52,59,60^ The spectra have strikingly similar cellulose NMR shifts such that the six major glucose environments a, b, c, f, g, j can be seen in a wide range of plants. This is illustrated in Fig. 3 using a comparison of the 30 ms ^13^C CP MAS PDSD NMR spectra of a eudicot (poplar), a monocot grass (*Brachypodium*) and a gymnosperm (spruce) in the C4-C1 and the C6-C4 regions. These six major glucose environments identified in poplar are always present in the other plants, but the relative amount of each environment varies. For example, spruce wood appears to have relatively more of site b versus a or c whereas in poplar wood cellulose these are similar in quantity. Regarding the minor glucose environments, *Brachypodium* has significantly more of site d compared to both poplar and spruce wood. The environment e as observed in primary cell walls^54^, whilst not visible in Fig. 3, is very minor in all three plants. There is one additional environment s in domain 1 of the spectra of spruce which we speculate could arise due to a different microfibril habit or cellulose interactions with galactoglucomannan, the major hemicellulose in softwoods. There are some small differences in the shifts for some of the minor glucose environments between plants. For example, we found that the minor environment d is generally broader than sites a, b, and c with its C4 ^13^C NMR chemical shift varying by ∼ 0.3 ppm indicating that there are some differences in the local environment of site d. In summary, the six major glucose environments (a, b, c, f, g and j) can be seen in the cellulose microfibrils of all the plants we have studied, reflecting common aspects of the structure of the cellulose microfibrils. The differences in abundance of the glucose environments could arise from changes in microfibril size or shape as well as interactions with hemicelluloses. Therefore, identifying the location of each of these glucose environments within the microfibril will provide invaluable insight into the different cellulose structures observed across various plant cell walls.

**Figure 3:**
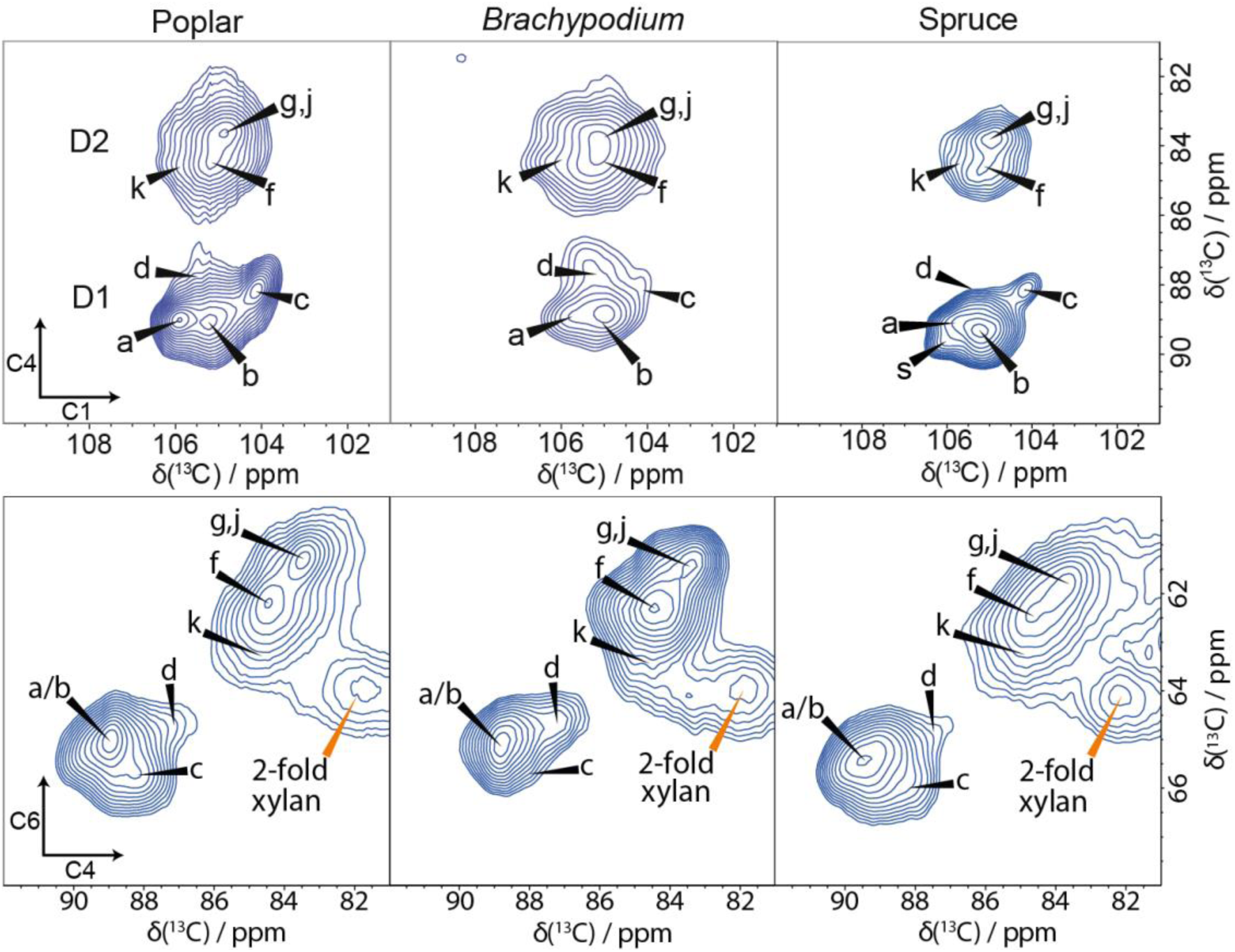
**Glucose environments in cellulose are common across different plants**. A comparison of the C4-C1 and C6-C4 regions of 30 ms ^13^C 2D CP MAS PDSD NMR spectra of poplar wood (a eudicot), Brachypodium mature leaves (a monocot) and spruce wood (a gymnosperm). Common glucose environments are labelled. Note that, whilst the relative amounts vary, the cellulose environments are common throughout. The spectra were recorded as in Fig. 1.

### 2.3. Xylanase-treated holo-cellulose nanofibrils (hCNFs) maintain the native plant cellulose structure of poplar wood

Whilst studying the native plant cellulose in-situ is ideal to ensure minimal disruption, the ^13^C MAS NMR spectrum can be crowded since signals from hemicelluloses and lignin are present within the same region which tends to complicate the analysis and limit the resolution of cellulose environments in the spectrum. The glucose environments in the microfibrils may also be influenced by interactions of surface residues with hemicellulose or lignin. By removing the lignin during preparation of holo-cellulose nanofibrils (hCNFs), we can maintain the cellulose microfibril structure whilst simplifying the ^13^C MAS NMR spectrum.^56^ TEM images, seen in SI Fig. 2, of the hCNFs sample show long and thin loosely bundled cellulose microfibrils of ∼3 nm in width. To reduce any effect on the spectrum of xylan presence, we removed the majority of xylan using xylanase hydrolysis. We compared the ^13^C MAS NMR spectra of poplar wood to those of the xylanase-treated hCNFs to ensure the major cellulose environments are maintained. The comparison of the 1D ^13^C CPMAS NMR spectra are shown in Fig. 4a. There is a distinct increase in the apparent resolution of the 1D spectrum for the hCNFs sample which is particularly evident in the C1 region and to a lesser extent in the C6 region. There is also a significant change in the total signal in the C4 region as well as the ratio of domain 1 and domain 2 cellulose peaks. This change is partly due to the removal of both lignin and hemicelluloses as well as, perhaps, the loss of a less ordered (broad) cellulose component that is not part of the cellulose microfibril. This more disordered component is removed during the production of the hCNFs, whilst the xylanase treatment of the hCNFs sample removes most of the xylan hemicellulose without affecting the ratio of the two cellulose domains as shown in SI Fig. 3.

**Figure 4:**
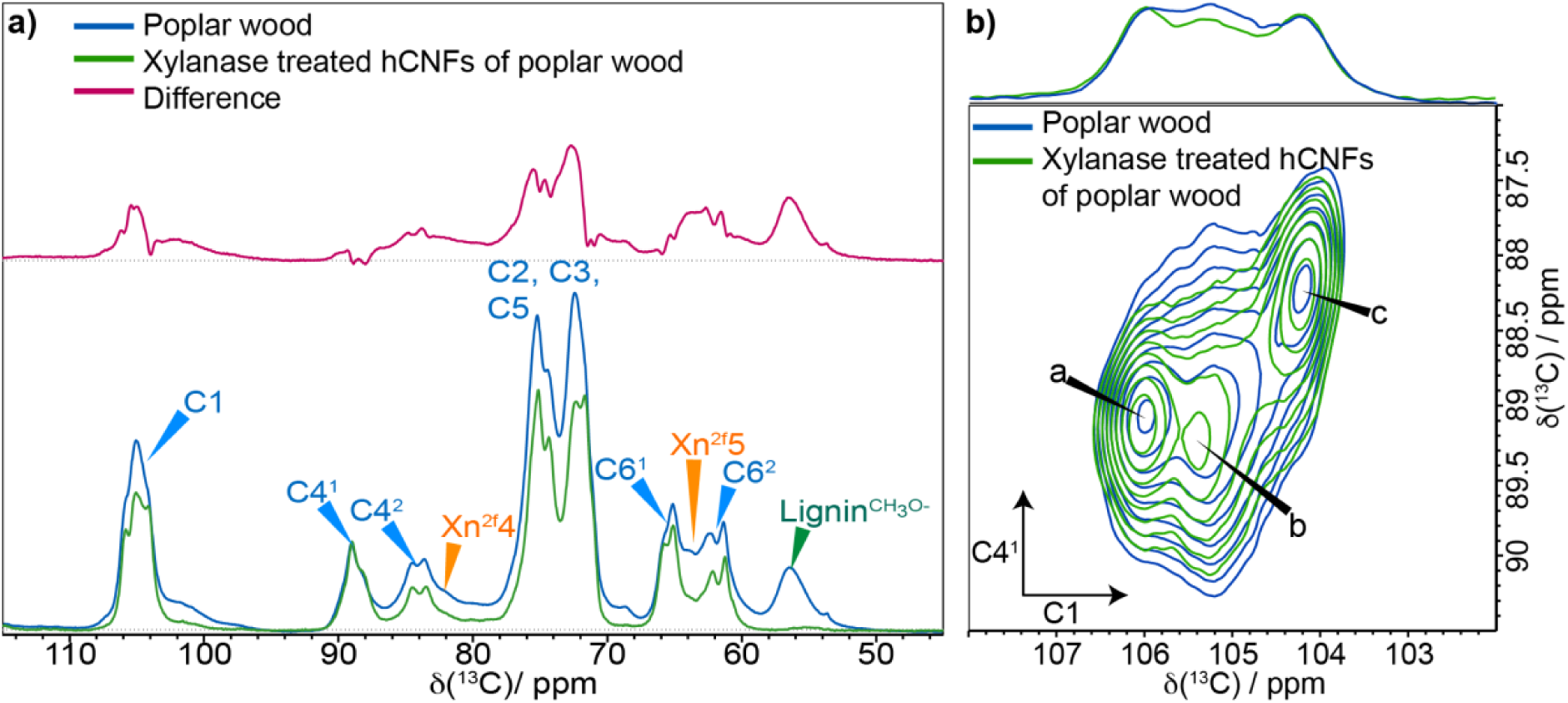
Comparison of poplar wood and xylanase-treated hCNFs of poplar wood. a) The neutral carbohydrate region of ^13^C 1D CP MAS NMR spectrum. The comparison (normalised to C4 at 89 ppm) shows the difference in the 1D CP spectrum of poplar wood (blue) and xylanase-treated hCNFs (green) when hemicellulose and lignin have been removed. It is clear the production of hCNFs results in the removal of most of a broad background signal as shown by the difference (red). **b) SCW cellulose of poplar wood maintains its structure in the fibrillation process to form hCNFs.** Comparison of the cellulose C4-C1 domain 1 region of 30 ms ^13^C 2D CP PDSD MAS NMR spectra of poplar wood (blue) and xylanase-treated hCNFs of poplar wood (green). Sum projections of the C4-C1 domain 1 region are shown at the top. Spectra were recorded as in Fig.1.

A comparison of 2D 30 ms ^13^C CPMAS PDSD NMR spectra of poplar wood and xylanase-treated hCNFs for the C4^1^-C1 and C6-C1 regions, shown in Fig. 4b and SI Fig. 4 respectively, confirms that the domain 1 glucose environments a, b and c remain almost completely unchanged by the production of hCNFs from poplar wood. The domain 2 environments show some slight ^13^C NMR chemical shift changes, typically < 0.3 ppm, in both the f and g/j environments. As the domain 2 environments are surface chains of the microfibril, this could be due to changes in their interactions caused by the removal of both lignin and some xylan. Since there were no substantial changes in the ^13^C NMR chemical shifts, we have determined that the production of the hCNFs causes relatively minimal disturbance to the cellulose microfibril structure. Therefore, these isolated fibrils can be used to help identify the six major glucose environments within the native cellulose microfibril. Interestingly, we find that the line widths of the major environments in spectral domain 1 and domain 2 are similar as illustrated in SI Fig. 5 which shows the C6 region of the CP refocussed INADEQUATE spectrum of the xylanase- treated hCNFs. The similar line widths for the major environments, summarised in SI Table 1, indicates that these different glucose sites in both these spectral domains have a similar local order.

**Figure 5:**
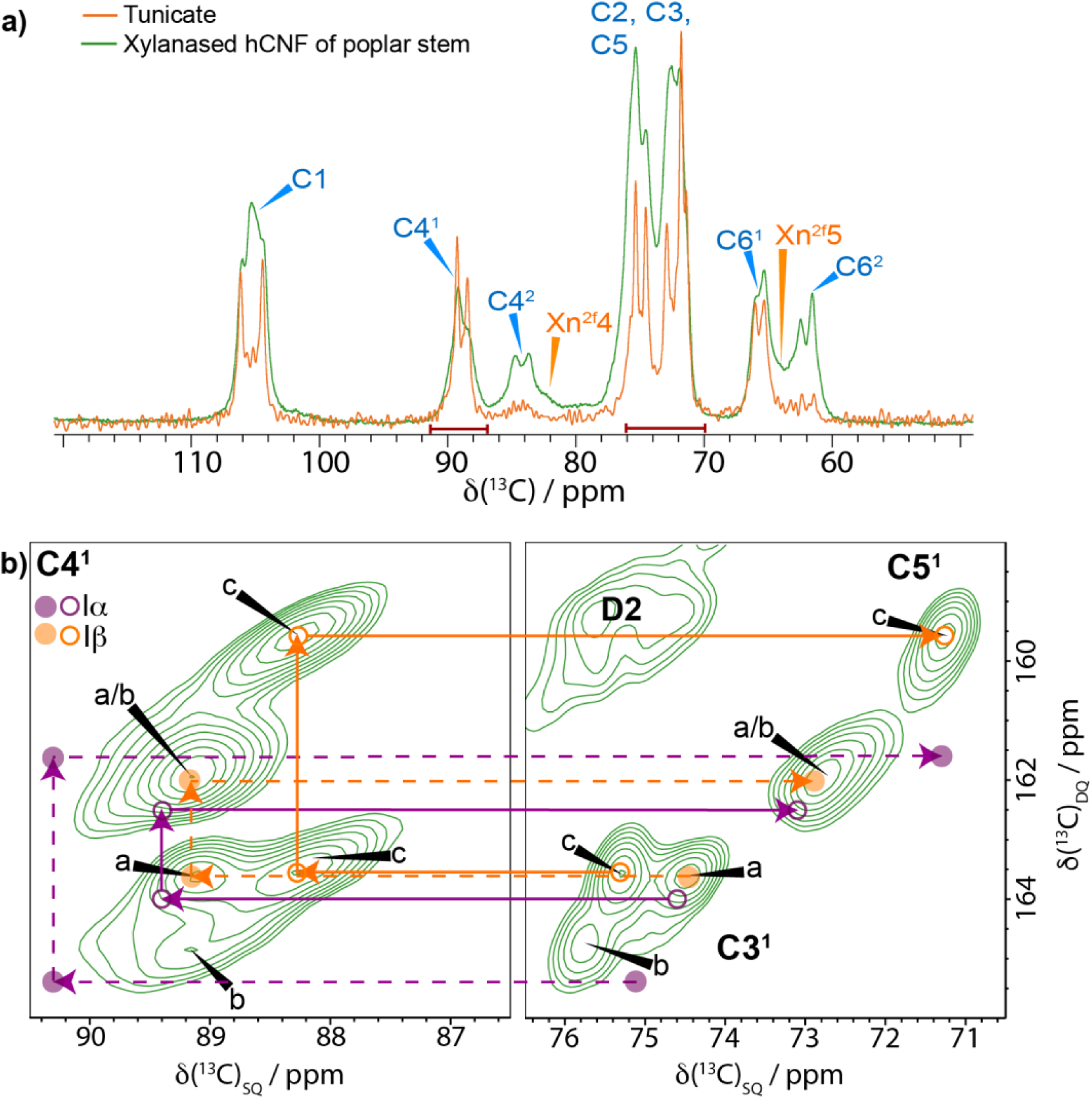
Plant secondary cell wall cellulose has a cellulose Iβ component but no cellulose Iα component. **a)** Comparison of 1D ^13^C CP MAS NMR spectra of tunicate (orange) and xylanase- treated hCNFs of poplar stem (green). **b)** Comparison of the domain 1 C4-C3/C5 region of a ^13^C 2D CP refocussed INADEQUATE MAS NMR spectrum of xylanase-treated hCNFs of poplar wood with the ^13^C chemical shifts of cellulose Iα and Iβ, as given by Brouwer and Mikolajewski^23,57^ (see SI Table 1). The filled and empty circles correspond to two distinct ^13^C chemical shifts in the asymmetric unit cells of cellulose Iα and Iβ.^23,57,64^ The Iβ cellulose shifts are within error (± 0.1 ppm) identical to those of cellulose sites a and c of xylanase-treated hCNFs of poplar stem. The Iα cellulose positions do not match the ^13^C NMR chemical shifts of any glucose environment acquired from the native never-dried plant cell wall. Spectra were recorded as in Fig. 1.

### 2.4. Plant cellulose microfibril core environments are identical to tunicate cellulose Iβ

The improvement in the resolution of ^13^C MAS NMR spectra makes analysis of the xylanase- treated hCNFs from poplar wood ideal for assigning the distinct ^13^C NMR chemical shifts to specific locations within the microfibrils. We compared the ^13^C NMR chemical shifts of our xylanase-treated hCNFs to those of a tunicate (cellulose Iβ) sample and to the ^13^C chemical shift assignments of cellulose I of Brouwer et. al. ^23^ In Fig. 5a, a comparison of the 1D CP MAS NMR spectra of tunicate (cellulose Iβ) and the xylanase-treated hCNFs sample shows that the tunicate shifts precisely overlay a subset of the peaks in the hCNFs spectrum. Indeed, the ^13^C chemical shifts for the C1 of a and c match the C1 ^13^C chemical shifts of those for tunicate cellulose. We next compared our ^13^C chemical shift assignments from the high-resolution ^13^C 2D CP MAS refocussed INADEQUATE NMR spectrum of xylanase-treated hCNFs with the ^13^C NMR chemical shifts of both cellulose Iα and Iβ. None of the domain 2 surface glucose environments ^13^C NMR chemical shifts are close to those in either of these cellulose allomorphs (SI Table 2). This means we only need to consider the ^13^C chemical shifts of spectral domain 1 glucose residues, where there are only 3 major sites: a, b and c.

Fig. 5b shows the C4-C3/C5 region of a ^13^C CP MAS refocussed INADEQUATE NMR spectrum of the xylanase-treated hCNFs with the expected Iα and Iβ ^13^C chemical shift positions highlighted on the spectrum. Interestingly, the C3, C4 and C5 shifts of glucose environments a and c, as in the 1D comparison with tunicate, are near identical to those of cellulose Iβ. Indeed, all the ^13^C chemical shifts for sites a and c, apart from C1, are within ∼0.1 ppm of those reported for cellulose Iβ (SI Fig. 6 and SI Table 2).^23^ Since the only substantial difference we observe from the previously reported Iβ ^13^C chemical shifts is in C1, it seems likely that the Kono et al.^21,22^ assignment was incorrect. This is highlighted in SI Fig. 7, where it can be seen that with the swapping of the C1 assignments there is alignment with those observed in our spectra. This misassignment by Kono et al.^22^ presumably arose because the ^13^C CP MAS refocussed INADEQUATE NMR spectrum was used for their assignment and both glucose environments in cellulose Iβ have nearly identical C2 ^13^C NMR chemical shifts making it difficult to distinguish the associated C1 chemical shifts. In contrast by using the 30 ms CP MAS PDSD NMR spectrum (SI Fig. 7) together with the ^13^C CP MAS refocussed INADEQUATE NMR spectrum we could determine confidently the C1 ^13^C chemical shifts for environments a and c. Indeed, very recently Brouwer and Mikolajewski have also noted that the Kono et al. C1 assignments should be exchanged.^57,58^

**Figure 6:**
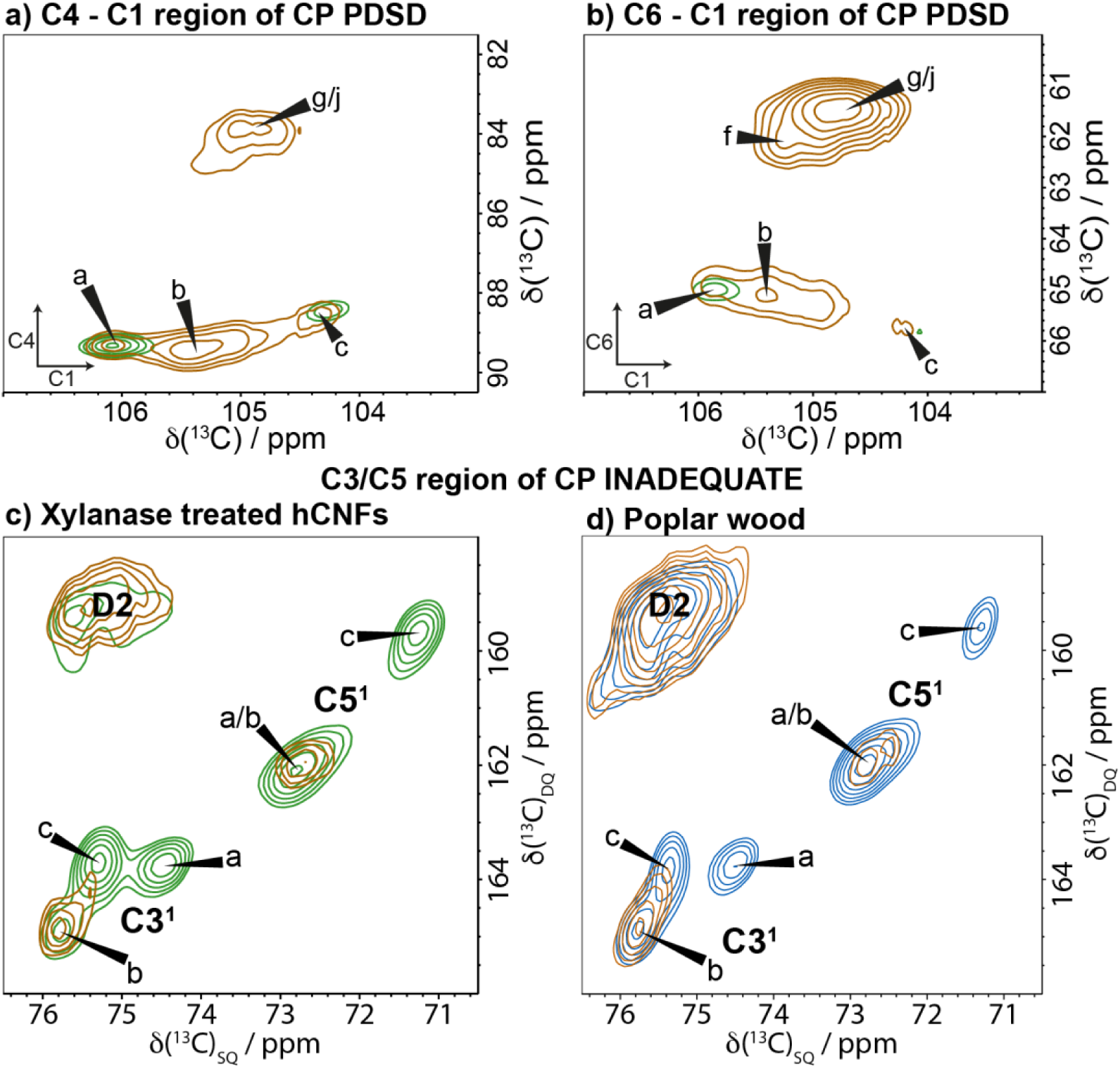
Domain 1 glucose environments in cellulose are not solely interior chains. Comparison of the standard (green) and water edited (brown) 30 ms ^13^C 2D CP MAS PDSD NMR spectra of xylanase-treated hCNFs of poplar wood. **a)** the C4-C1 region, **b)** the C6-C1 region. Spectra were normalised to glucose environment a. **c)** Comparison of the C3/C5 region of the standard (green) and water-edited (brown) CP refocussed INADEQUATE spectra of xylanase-treated poplar wood. **d)** Comparison of the C3/C5-C4 region of the standard (blue) and water-edited (brown) CP MAS refocussed INADEQUATE NMR spectra of poplar wood. Both c) and d) were normalised to glucose environment b. Spectra were recorded as in Fig. 1.

**Figure 7:**
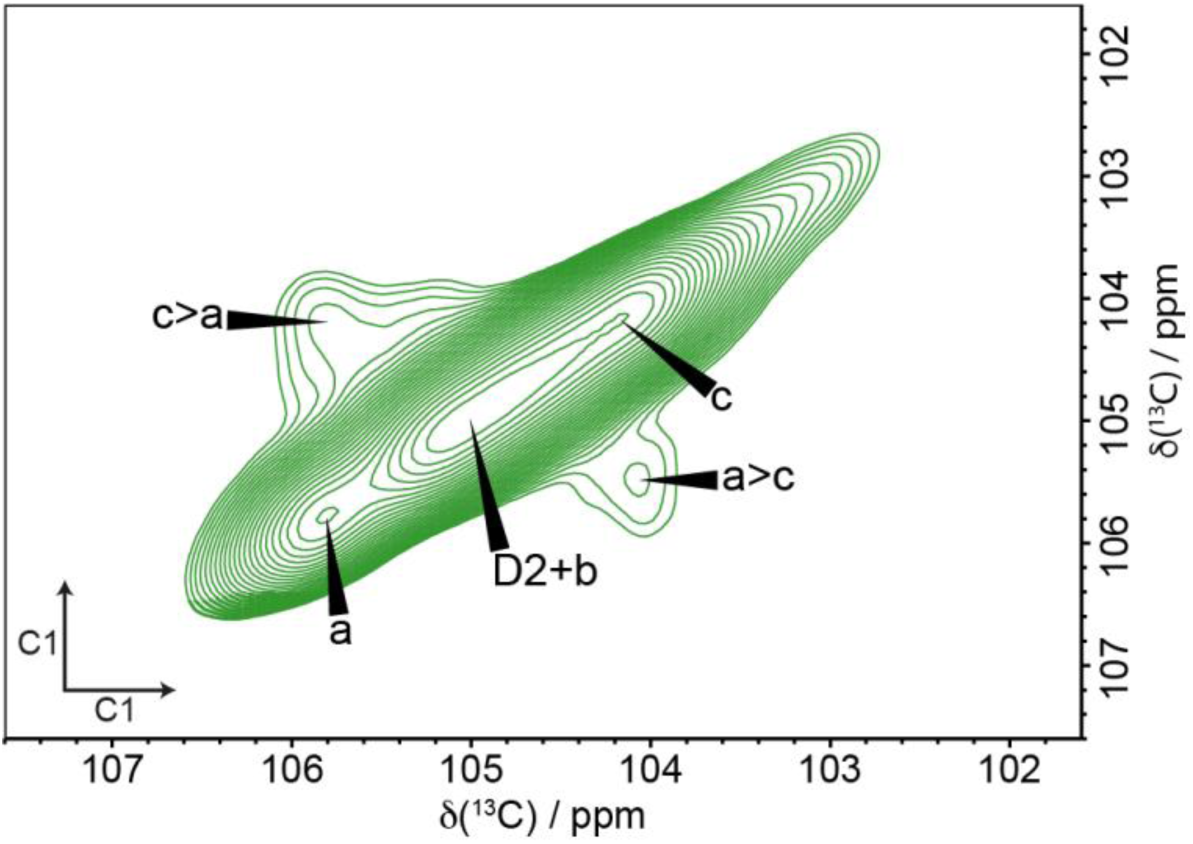
Domain 1 glucose environments a and c are in close proximity. The C1-C1 region of a 200 ms ^13^C 2D CP MAS PDSD NMR spectrum of xylanase-treated hCNFs of poplar wood shows cross-peaks between a and c. Spectra were recorded as in Fig. 1.

With this new NMR shift assignment of cellulose Iβ all the shifts of microfibril core glucose environments a and c match closely to those for tunicate cellulose Iβ. Furthermore, we can now assign environment a as origin chain and environment c as centre chain, since DFT calculations for cellulose Iβ predict the C1 chemical shift of the residues in the origin chains to be ∼2 ppm higher than the C1 of residues in the centre chains, as seen here for environments a and c respectively.^54,65^ We next considered evidence for the presence of cellulose Iα. Despite environment b having similar ^13^C chemical shifts to those of one of the Iα glucose environments for both C1 and C4 (SI Table 2), there is no sign of the second Iα glucose environment with a C4 at 90.3 ppm which would be in similar quantities to site b. It is evident from Fig. 5 that the shifts of cellulose Iα are very different from those observed for xylanase- treated poplar hCNFs. Since this is true for all the plant samples analysed here, there is a clear absence of any cellulose Iα in plants. This observation is contrary to the long-held belief that plant cellulose is a mixture composed of largely cellulose Iβ with varying amounts of the cellulose Iα allomorph ^26,30^ whereas, in fact, it is solely a cellulose Iβ structure.

Having determined that two of the main spectral domain 1 environments correspond to centre and origin chains in the classical cellulose Iβ structure, we wanted to understand the nature of the third, remaining, domain 1 environment, b. Since site b is from domain 1, it has been assumed to be interior to the microfibril. However, given this is not a cellulose Iα nor Iβ environment, its position within the microfibril is unclear. To investigate this further, two 2D water-edited NMR experiments were undertaken to probe the glucose environments that are closest to water, specifically (i) a 30 ms CP MAS PDSD and (ii) a refocussed CP MAS INADEQUATE experiment. Fig. 6a and 6b shows the C4-C1 and C6-C1 regions of a water edited PDSD spectrum compared with a standard 30 ms CP MAS PDSD spectrum (see SI Fig. 8). As expected for surface glucose residues, the signal for the domain 2 sites f, g and j is enhanced in both of these regions indicating that these residues are more water accessible than the microfibril core origin and centre chain sites a and c. Interestingly, Fig. 6a shows that the water edited C4-C1 peak for glucose environment b is also significantly enhanced in a similar way to the surface glucan sites in spectral domain 2, indicating that b is also closer to water than the microfibril core sites a and c. This observation suggests that glucose environment b of domain 1 is therefore, unexpectedly, a surface chain of the microfibril. Interestingly, for the C6-C1 region shown in Fig. 6b the signal from b is enhanced in the water edited experiment over the signal from the core a and c environments, but to a lesser extent than for the domain 2 surface glucose environments. Being in domain 1, environment b likely reflects a glucose residue with the C6 hydroxymethyl in a *tg* conformation. This conformation may arise because the C6 is facing toward the interior of the cellulose microfibril where the hydroxymethyl group hydrogen bonds with other glucose residues.^16^ Hence, in the glucose residue of environment b, C6 is further from water than the C4 and further from water than the other surface environments where the C6 has a water-facing *gt* or *gg* conformation. The C3/C5 region of the water edited ^13^C CP MAS refocussed INADEQUATE NMR spectrum of both the xylanase- treated hCNFs (Fig. 6c) and of poplar wood (Fig. 6d) also shows that for both samples site b is in closer proximity to water than the core glucose environments a and c. This proximity difference is observed for both poplar wood and the hCNFs sample indicating that it is not an artefact of the hCNFs sample and is representative of the native cellulose microfibril. Additionally, the C6 region of the ^13^C CP MAS refocussed INADEQUATE NMR spectrum shows that all three major domain 2 glucose environments are close to water with perhaps site g being marginally closer compared to sites f and j (SI Fig. 9). In summary, the water proximity experiments reveal that, in contrast to the widely accepted view^32,43,66^, not all the domain 1 glucose residues are in the interior of the microfibril.

**Figure 8:**
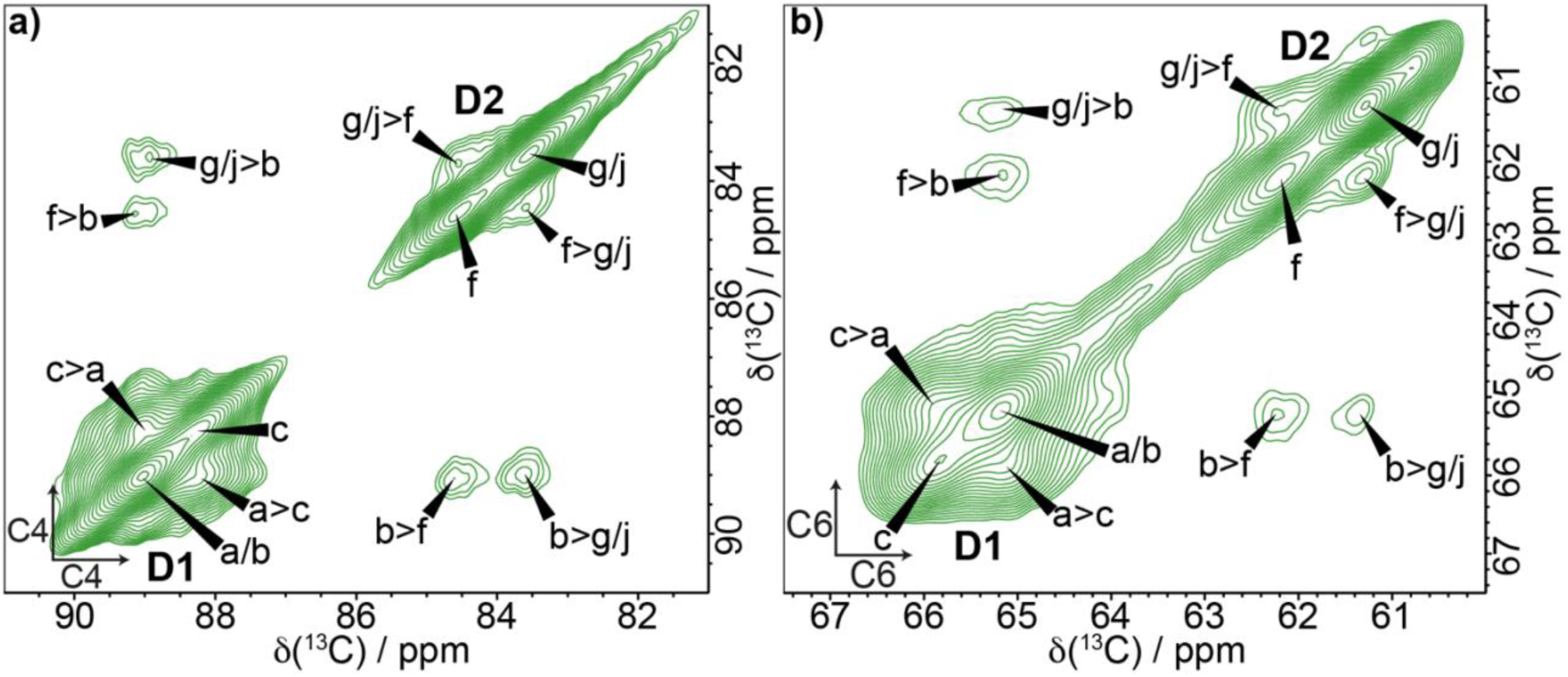
**Domain 1 glucose environment b and the domain 2 glucose environments are in close proximity**. A 400 ms ^13^C 2D CP MAS PDSD NMR spectrum of xylanase-treated hCNFs of poplar wood. **a)** the C4-C4 region, **b)** the C6-C6 region. Spectra were recorded as in Fig. 1.

**Figure 9:**
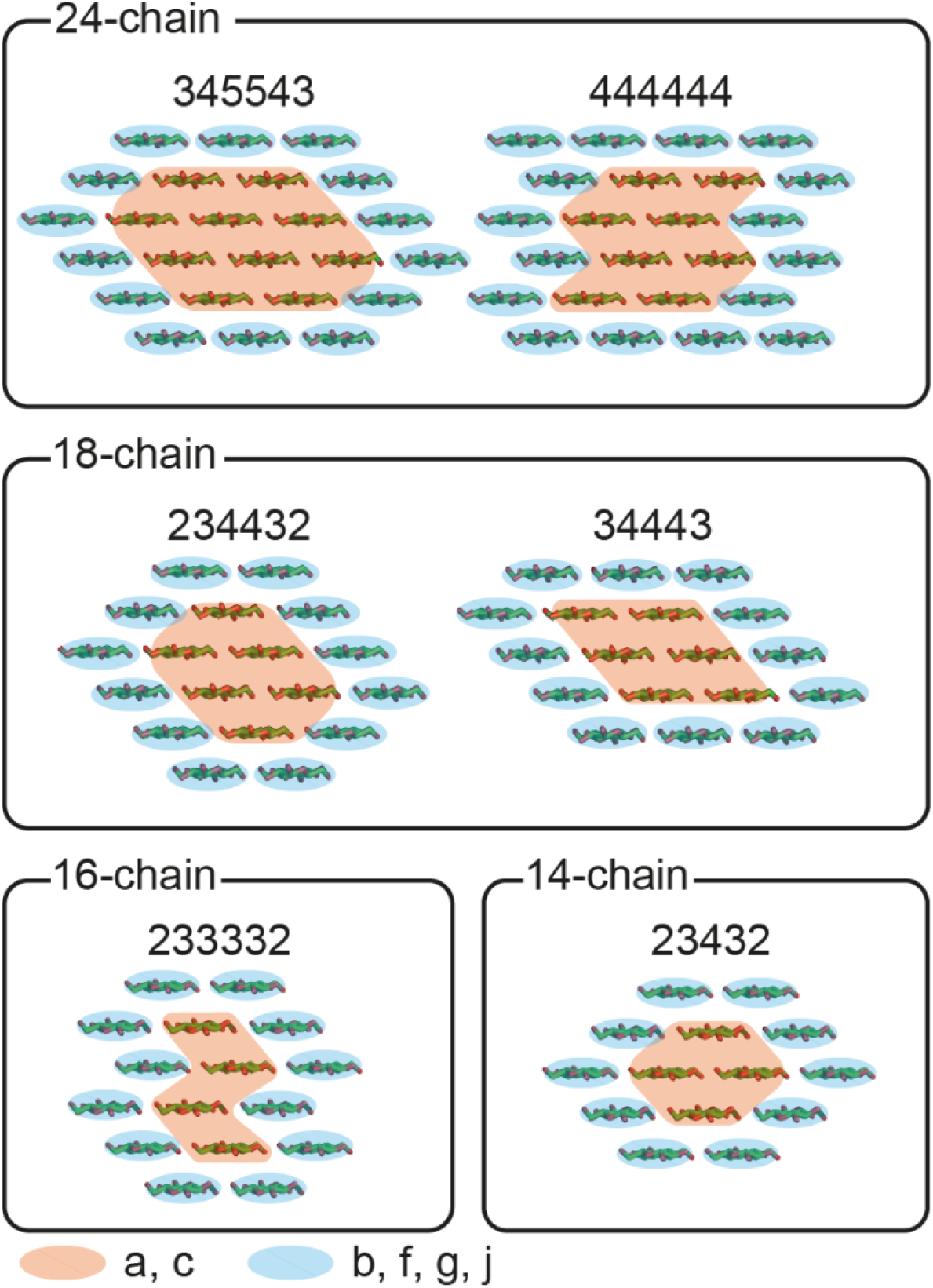
**Possible cellulose fibril habits that fit the interior to surface ratio of 1:2 as determined here by ^13^C MAS NMR for the xylanase-treated hCNFs of poplar stem.**

To investigate the relative proximities of the different glucose environments within the microfibril, two longer (200 ms and 400 ms) mixing time ^13^C CP MAS PDSD NMR experiments were performed. As these longer mixing time experiments are probing distances up to ∼ 5-8 Å, additional cross-peaks between glucose residues in adjacent sites within individual fibrils are observed. Fig. 7 shows that at a mixing time of 200 ms the C1-C1 region of the ^13^C CP MAS PDSD NMR spectrum gives clear cross peaks between a and c only. This shows that the centre and origin chain environments a and c are closer to each other than either are on average to residue environment b. This is consistent with our finding that b is a surface environment and is not situated with a and c in the core of the cellulose microfibril. The first clear cross peaks between glucose residues in the two different domains can be seen in the C4-C4 region of the 400 ms ^13^C CP MAS PDSD NMR spectrum (Fig. 8a). There are cross peaks from the surface domain 2 glucose environments f and g/j to the single C4^1^ peak corresponding to environments a and b. Given we know that environments a and c are particularly close to each other, and there are no cross peaks observed from domain 2 to site c, the cross peak to domain 2 therefore comes from environment b only. This proximity of b and not c to domain 2 glucose residues is also observed in the C6-C6 region of the 400 ms CP MAS PDSD spectrum (Fig. 8b). There are also cross peaks between the different domain 2 surface environments (f and g/j) of cellulose, consistent with them all being located on surfaces. The proximities of b to the domain 2 environments provides further confirmation that b is one of the four major surface glucose residue environments.

Given the new assignment of the core (sites a and c) and surface environments (b, f, g, j), the true ratio of interior vs surface of the microfibril can be estimated by integrating the C1 region of the CP refocussed INADEQUATE spectrum where core sites a and c are distinguishable from the surface glucose environments (b + D2) (see SI Fig. 10a). Since the refocussed INADEQUATE spectrum is not necessarily quantitative, the reliability of this integration was confirmed by comparing 1D CP, quantitative DP and 1D double quantum (DQ) filtered spectra for the xylanase-treated poplar hCNFs sample. Two different echo times were used in the 1D DQ filtered experiment to determine whether this affected the relative amplitude of the peaks. When normalised to the core C1 peak (site a) at 106.1 ppm there was very little change in the relative intensities of different glucose environments within the C1 region as shown in SI Fig. 10b. Hence, the integration of the C1 area of the CP refocussed INADEQUATE (see SI Fig 10a) can be used to give an estimate of the relative amount of interior sites a and c versus surface sites b +D2 (f, g, j) where we find a value close to 0.5, i.e. an interior: surface (i:s) ratio of 1:2.

## 3. Discussion

The structure of native plant cellulose has been studied by a wide range of techniques for decades.^25,37,67–69^ However, despite this, the microfibril structure remains to be fully elucidated with its core structure, microfibril size and habit still widely disputed.^9,11,15^ In this work we have studied a range of samples and have used two-dimensional high magnetic field MAS NMR of never-dried cell walls to resolve some of the misinterpretations and inconsistencies that have arisen partly due to the overlap of signals from the cell wall components using 1D NMR. We have taken several steps to achieve the high resolution observed in the spectra we acquired. We only used never-dried cell walls, because it is observed in softwoods^53^ and hardwoods^36^ that dry plant samples give significantly broader NMR spectra. We have also acquired for sufficiently long in the indirect dimension alongside utilising some of the highest magnetic fields available to achieve higher resolution. The use of the hCNFs in this work also increased the apparent resolution of the cellulose spectrum, in part by removing components such as lignin and hemicelluloses like xylan that might alter cellulose chemical shifts.

The breadth of studies regarding cellulose crystallinity across plant and material sciences has given rise to terminology which can be confusing, especially when the same terminology is applied to data acquired by different methods. For example, describing cellulose as crystalline or amorphous has different meanings in NMR studies compared to diffraction experiments. Measuring crystallinity of a cellulose sample by NMR is based on a much shorter range order than that probed by diffraction techniques.^70^ Historically the glucose residues observed in the two domains of NMR spectra were assigned as crystalline and amorphous cellulose because the characterised crystalline forms of cellulose I have ^13^C chemical shifts in the spectral domain 1 (C4∼89 ppm) region.^44,71,72^ Surface residues that give peaks in spectral domain 2 are sometimes described as paracrystalline or amorphous^27^ leading to the ratio of the C4^1^ and C4^2^ peaks from the 1D NMR spectrum being commonly called (predominantly by material scientists) the crystallinity index (CI). This index is currently routinely used to describe the crystallinity of cellulosic samples alongside crystallinity measured by x-ray diffraction.^44,71^ The measured values of crystallinity by NMR versus diffraction techniques are always significantly different, although the general trends across a range of samples tended to be consistent.^44^

In our work, we find that the line widths of the signals from the six glucose environments within the surface and core of the microfibril are very similar: e.g., the C6 region of the CP refocussed INADEQUATE spectrum of the xylanase-treated hCNFs shown in SI Fig. 5. SI Table 1 summarises the observed linewidths. NMR chemical shifts are sensitive only to short range structure (< ∼ 5 Å), and so the similar narrow linewidths indicate that the glucose environments contributing to both domains have similarly high short-range order. Therefore, the microfibril core and surface glucose residues cannot be divided simply into crystalline and amorphous cellulose on an NMR scale. There is, however, a weak (<10%) broad signal in the 1D CPMAS spectrum of poplar stem seen in Fig. 4a underlying the domain 2 region that is removed in the production of the xylanase-treated hCNFs sample. Some of this less ordered component is likely to be lignin as well as xylan but it could also include a more disordered portion of cellulose that is known to be easily hydrolysed.^73^ We note that this disordered material in the intact wall is lost during isolation of the hCNFs, calling into question the belief that microfibrils have substantial regions of amorphous cellulose interspersed along each fibril between crystalline domains.^17^ Therefore, whilst there might be a small component of less ordered cellulose giving rise to signal in the domain 2 region in cell walls and processed lignocellulose, the term amorphous cellulose does not accurately describe the origin of this NMR signal when referring to cellulose microfibrils. We further note that, depending on the sample, the ratio of spectral domain 1 to domain 2 signals could be influenced by the microfibril dimension, as well as the presence of any signal from other cell wall components which does not necessarily arise from cellulose (e.g. lignin, hemicelluloses). Hence the traditional CI as measured by NMR is a misnomer; it does not measure cellulose crystallinity.

Some groups have designated all spectral domain 1 glucose environments as arising from the interior of the microfibril with a structure that is a mixture of the two cellulose I allomorphs or even proposing a distinct structure.^15,48,50^ We emphasise that high resolution MAS NMR spectra as presented in this work have been needed to resolve signals from these domain 1 residues to substantially test these hypotheses. The ^13^C NMR chemical shift is very sensitive to the local environment allowing us to distinguish differing conformations, such as 2-fold and 3-fold xylan structures^34^ as well as changes in hydrogen bonding. The 2D solid-state NMR spectra of several samples presented here has allowed distinct and recurring glucose residues in the cellulose microfibrils to be resolved and ^13^C NMR chemical shifts to be fully assigned. In contrast to previous work on primary cell walls^54^, we found that native plant cellulose has just six major glucose environments, with only three of these contributing to the spectral domain 1 region. Within domain 1 we have now unambiguously assigned the glucose environments a and c as cellulose Iβ since the ^13^C NMR shifts of our tunicate sample as well as those of cellulose Iβ given by Brouwer and Mikolajewski are identical to those found here.^57,58^ We confirm, using the 30 ms ^13^C CP MAS PDSD NMR spectra of our samples, that the C1 ^13^C chemical shift of the two glucose residues of tunicate cellulose Iβ had previously been misassigned by Kono, as was recently also suggested by Brouwer and Mikolajewski.^57,58^ With this assignment the differences in the two residues are now more consistent with DFT calculations of cellulose Iβ origin and centre chain environments,^65^ allowing us to assign origin chains to site a and centre chains to site c (SI Table 1). Note that, since these core environments have the ^13^C chemical shifts of cellulose Iβ, they must be surrounded by chains packed in a cellulose Iβ fashion, and hence we deduce that the microfibril is entirely cellulose Iβ.

This assignment of cellulose Iβ structure of the native plant cellulose microfibril is consistent with the clear lack of cellulose Iα signals in the 2D MAS NMR spectra. The assertion that native plant cellulose is a mixture of the two cellulose I allomorphs originally arose by comparing 1D NMR spectra of native plant cellulose with those of the two biological crystalline allomorphs of cellulose.^30^ It was found that there was an additional component in the domain 1 C4 region which meant that the origin and centre chains were not in a 1:1 ratio as expected of a cellulose Iβ crystal. By adding a small amount of cellulose Iα structure to the simulation of the 1D NMR spectra the resulting fit appeared good.^30,67^ However, if there were cellulose Iα present in plants, we would expect to observe two equal intensity Iα glucose environments contributing to the domain 1 region of the 2D spectra. On the contrary, we found that there is only one further major glucose environment, b (see Fig. 5b). This glucose environment b has similar C1 and C4 shifts to one of the glucose residues of cellulose Iα perhaps leading to the earlier misinterpretation of the 1D spectra. We have now conclusively shown that the microfibril is solely cellulose Iβ, and so cellulose modelling, computational studies and interpretation of FTIR and diffraction patterns should be based solely on this Iβ structure.

Glucose residues with ^13^C chemical shifts in domain 1 (C4 around 89 ppm) have long been considered to reside in the crystalline core of plant cellulose.^43,55^ However we have now shown, using water-edited experiments, that the third major glucose environment in spectral domain 1, site b, is a residue on the surface of the microfibril. We previously suggested^34^ that there could be a surface component in the domain 1 region and recently this view has been supported by the work of Addison et al.^36^ Calculations have shown that a change in C6 hydroxymethyl conformation from *tg*, to *gt/gg* changes the C4 ^13^C chemical shift by ∼ 5 ppm.^47,74^ Adoption of the *tg* conformation is likely to occur where the C6 is facing towards the inside of the microfibril where the orientation is fixed by hydrogen bonding to other glucosyl residues rather than water. Therefore, our experiments indicate that the glucose residue in environment b, is a surface glucan chain which has a *tg* conformation, perhaps due to the C6 carbon facing inwards towards other chains in the microfibril. The previously proposed models requiring multi-layered microfibril environments^75^ or bundling^43^ to generate additional interior chains are not necessary to explain the major glucose environments within spectral domain 1. It is now clearly more appropriate to describe spectral domain 1 glucose residues as possessing *tg* C6 hydroxymethyl conformation, and that spectral domain 1 reflects neither crystalline cellulose nor the exclusively interior of the microfibril.

The differing assignments of spectral domain 1 and spectral domain 2 has led to a range of different models to illustrate the structure of native plant cellulose which has been widely debated.^9,37,43,54,55^ Whilst some models are able to explain some of the phenomena observed in cellulose there is no overarching model that is able to describe cellulose over a wide range of materials. Previous studies have used the ratio of C4^1^ and C4^2^ from 1D NMR spectra as a measure of microfibril size, which has resulted in variable estimations such as 18-24 chains in a microfibril or alternatively microfibril bundling models.^9,43^ As we now know, domain 1 is does not arise solely from interior sites, thus the ratio of C4^1^ and C4^2^ will give an overestimate of the interior of the microfibril and thus overestimate the microfibril size. The ratio of the two domains could still be utilised for characterisation as it is loosely related to the volume of the microfibril, but care is needed in using this interpretation. In this work, we have provided a more accurate measure of the interior to surface ratio (i:s) for the microfibril of poplar wood which was found to be ∼0.5, i.e., i:s = 1:2. We also gain insight into the proportions of the environments in the microfibril, namely origin (a) and centre (c) chains are similar in proportion (1:1 ratio) as well as, for xylanase-treated poplar wood microfibrils, the relative amount of surface environment b is less than sites a or c. Given the constraints on the microfibril dimensions from biophysical measurements the upper limit is a microfibril with around 18-24 chains, however there are several habits that could fulfil these criteria.

It should also be noted that NMR is a bulk technique and the spectra and quantification of core to surface is the result of an average of many microfibrils, so care is needed in the interpretation. However, one could assume that most common microfibrils present in the material will dominate the spectra. The ratio of i:s = 1:2 is consistent with an 18-chain microfibril with a habit that has 6 interior chains and 12 surface chains, but it could also be consistent with a 24-chain microfibril with 8 interior and 16 surface chains. Thus, for the xylanase-treated hCNFs of poplar wood, this would mean for a slice through an 18-chain microfibril that there are three of each site a and c chains and hence potentially 2 chains of site b (since b < a and c see Fig. 5) with the remaining 10 chains being domain 2 environments (f, g and j). The ratio of the surface and interior ratio alone is not sufficient to give a value for the size of the microfibril. As shown in Fig. 9, there are several different sized microfibrils that could have a ratio close to 1:2. Hence further understanding of the relative proportions of the different glucose environments reduces the possible cellulose microfibril habits. For example, two potential 18-chain habits 234432 and 34443 have the observed interior to surface ratio of 0.5. However, given that the cellulose Iβ structure has origin (a) and centre (c) chains in alternating sheets, the 34443 with three core layers may not have equal amounts of sites a and c. On the other hand, the habit 234432 would fit this constraint.

Whilst we cannot yet propose a definitive microfibril structure based on the NMR data, when we understand where the surface glucose NMR environments reside in the hydrophilic and hydrophobic surfaces of the cellulose microfibril, their relative proportions will provide a fingerprint of the cellulose microfibril structure. The relative amount of surface site b compared to core chain sites a and c for isolated microfibrils is likely to vary with any changes in the microfibril size, habit, and hemicellulose interactions in different plants. Therefore, the relative proportions of the six major glucose residue environments will be an important diagnostic tool for discovering cellulose variability, studying cellulose interactions, and for detecting changes during degradation or industrial processing of wood and other biomass. For example, in a study of cotton by Kirui et al.^37^ it was found that there is relatively more of the core glucose environments a and c compared to the surface environments (b, f, g, and j) which is consistent with cotton having a significantly larger fibril structure.^76^ On the other hand, we find the relative amounts of site a and c are less than site b in the *Brachypodium* sample we have studied in this work (Fig. 3) suggesting that there is either a smaller microfibril or different habit with fewer core chains compared to those proposed for poplar wood.

## 4. Conclusion

We have clarified that plant cellulose microfibrils have the cellulose Iβ structure; we do not detect any cellulose Iα. Moreover, we find no evidence for amorphous cellulose in the microfibrils with the surface glucose residues being well ordered. We show that a spectral domain 1 environment (site b), previously thought to be interior is from a surface glucose residue. This work changes widely held interpretations of solid-state ^13^C MAS NMR spectra since previous interpretations of sample crystallinity and surface to core ratios are shown to be flawed. The proportion of spectral domain 1 to domain 2 is better understood to reflect the ratio of C6 hydroxymethyl group conformations *tg* vs *gt/gg*, and so are influenced largely by whether the residue is hydrogen bonded to another cellulose chain or to water. Our results now also provide a strong basis for understanding cellulose microfibril surfaces as we have shown that there are four major types of surface residues which likely reflect, in a manner to be determined, the different hydrophobic and hydrophilic surfaces of microfibrils. The exciting prospect now arises of a more complete description and understanding of cellulose microfibril surfaces and their interactions and variations in both biomaterials and plant cell walls.

## Methods

### Sample Production and Preparation

The poplar stem *Populus tremula* × *tremuloides* used in this work was grown from poplar shoot cuttings which were allowed to root for 2 weeks before being transferred into the growth chamber. Here the saplings were grown in a ^13^CO_2_ atmosphere^34,77^ for ∼ 3 months to provide ∼97% ^13^C labelled material that is predominantly secondary cell wall. Only wood of lower (woody) never-dried stems was used after removing bark and cambium. The growth of spruce and *Brachypodium* was as described in Terrett et al. and Tryfona et al. respectively.^52,59^

To prepare hCNFs, the never-dried wood after removing bark and cambium was first delignified four times with 3% peracetic acid (PAA; 0.35 g pure PAA/g dry material, pH 4.8) at 85 °C for 45 minutes without stirring. Between cycles, used PAA was decanted, one water wash was done, and new 3% PAA was added. Delignified material (holocellulose) was washed with water after the fourth cycle until the conductivity was below 10 S/cm. The holocellulose was blended for two minutes to form a homogenous slurry. Holocellulose dispersion (0.1 wt%) was blended for 30 minutes in a Vitamix A3500i blender (Vitamix, US) to make a polydispersion. hCNFs were recovered as supernatant from the polydispersion by centrifugation at 5000 rpm for 15 minutes. hCNFs were concentrated by filtration onto a PVDF membrane and the wet cake was then freeze-dried and rewetted into the NMR rotor. The hCNF suspensions were digested with xylanases as described previously.^56^ The xylanase- treated hCNFs were then freeze-dried and rewetted before being packed directly into the NMR rotor.

The never-dried ^13^C enriched poplar wood was debarked and cut into small pieces of ∼ 1-2 mm size with a razor blade and then packed into a 3.2 mm MAS zirconia NMR rotor whilst removing excess water.

Tunicate (*Halocynthia roretzi*) was purchased at a local supermarket (Yoshiike, Japan). The cellulose was purified according to the method described previously.^78,79^ After the entrails were removed, the tunicate mantle was deproteinized and bleached by four cycles of alkaline treatment with 5% KOH (Fujifilm Wako, Japan) at 37℃ and oxidation with sodium chlorite (Fujifilm Wako, Japan) at 37℃ and pH 5, respectively. The bleached tunicate mantle was homogenized with a Vitamix V1200i blender (Vitamix, US) for 2 min at the top speed and was subsequently acid hydrolyzed with 50% sulphuric acid at 40℃ for 8 hours. The sediment was collected by filtration and thoroughly washed with deionized distilled water, followed by homogenization to gain an aqueous suspension of cellulose microcrystals. For ^13^C NMR, tunicate cellulose was centrifuged at 20,000g for 30 min to provide a more concentrated cellulose sample.

### TEM

A ThermoFisher Scientific (FEI) TalosF200X G2 microscope operating in scanning mode at 200 kV was used to obtain the TEM images.

### Solid-state NMR

Solid-state NMR experiments of poplar wood and the xylanase-treated hCNFs were acquired on a Bruker 1 GHz AVANCE NEO solid-state NMR spectrometer operating at ^1^H and ^13^C Larmor frequencies of 1000.4 and 251.6 MHz respectively using a 3.2 mm E^Free^ triple resonance MAS probe. All experiments were conducted at an input gas temperature of 10°C and an MAS frequency of 12.5 kHz with a recycle delay of 2 s. The ^13^C chemical shifts were determined

using the carbonyl peak at 177.87 ppm of L-alanine as an external reference with respect to tetramethylsilane. This referencing was confirmed to correspond with referencing to the adamantane CH_2_ peak to 38.48 ppm^80^ to ensure direct comparison with Brouwer and Mikolajewski.^23,58^ The 90° pulse lengths were typically 3.2 µs (^1^H) and 4.0 µs (^13^C). Cross polarization from ^1^H to ^13^C was achieved using ramped (70–100%) ^1^H radiofrequency amplitude^81^ and a contact time of 1 ms. SPINAL-64 decoupling was applied at a ^1^H nutation frequency of 70-80 kHz during acquisition and during the indirect dimension of 2D experiments.^82^ Sign discrimination in the indirect dimension of the 2D experiments was achieved using the States-TPPI method. For the principal assignments of cellulose environments, a CP refocused INADEQUATE ^13^C double-quantum (DQ) ^13^C single-quantum (SQ) correlation experiment was used whereby the evolution of DQ coherence for directly bonded carbons within the same glucan ring is correlated with directly observed SQ coherence.^63,83^ The acquisition time in the indirect dimension was 6.67 ms with a spectral width of 37.5 kHz and 128 acquisitions per *t*1 FID. The echo (τ-π-τ) duration, τ, was 2.24 ms giving a total echo time of 8.96 ms. Both intra- and intermolecular contacts were probed using 2D ^13^C-^13^C CP MAS proton driven spin diffusion (PDSD) experiments with mixing times of 30 to 400 ms.^84^ The acquisition time in the indirect dimension (*t*_1_) of the CP MAS PDSD experiments was 5.5–8.1 ms. The spectral width in the indirect dimension was 37.5 kHz with at least 64 acquisitions per *t*_1_ FID. The proximity of water to different cellulose environments was probed using both a water-edited ^13^C-^13^C 30 ms CP MAS PDSD experiment and water-edited CP MAS refocussed INADEQUATE experiment. These water-edited experiments are based on the normal CP MAS refocussed INADEQUATE and CP MAS PDSD experiment, however before the CP contact time there is a ^1^H *T*_2_ filter (spin echo) to remove signal arising from directly attached protons followed by a delay to allow for spin diffusion from the water protons to its near neighbours.^85^ The CP parameters were as stated above. The acquisition time for the water-edited ^13^C-^13^C 30 ms CP MAS PDSD experiment in the indirect dimension was 4.8 ms with a spectral width of 37.5 kHz and 576 acquisitions per *t*_1_ FID. The total ^1^H *T*_2_ filter was 320 µs followed by a spin diffusion delay of 2 ms. For the water-edited CP refocused INADEQUATE the acquisition time in the indirect dimension was 4.7 ms with a spectral width of 28.5 kHz and 960 acquisitions per *t*_1_ FID. The total ^1^H *T*_2_ filter was 280 µs followed by a diffusion delay of 2 ms. All 2D spectra were processed with Fourier transformation into 8 K (*F*_2_) × 2 K (*F*_1_) points with exponential line broadening of 20-50 Hz in *F*_2_ and cubed sine bell processing in *F*_1_ using Bruker Topspin v.3.6. Contour levels are x 1.1 throughout. The minimum contour is chosen to show the desired features. Further experimental details are summarised in SI Table 3.

## Data availability

Unprocessed NMR data files will be available from http://wrap.warwick.ac.uk

## Author contributions

AEP grew and harvested the ^13^C-labelled plants. PKD prepared the hCNFs, xylanase-treated hCNFs and imaged them, and TK sourced and prepared the tunicate sample. RC and RD conducted the NMR experiments and analysis. YY generated models of cellulose microfibril habits for conceptual analysis of glucose environments. RC, PKD, YY, TK, AEP, SPB, RD, and PD contributed to NMR data interpretation. PD and RD conceived and supervised the work. SPB and WTF contributed to experimental method development. RC, RD and PD wrote the paper with comments from all authors.

## Acknowledgements

This paper is dedicated to Ray Dupree, who enjoyed applying solid state NMR to address questions of cellulose structure, and who sadly died just before submission. Many thanks to Eva Hellmann from the Sainsbury lab at University of Cambridge for donating the poplar saplings that were used to grow the ^13^C material used in this work. Many thanks to Theodora Tryfona for providing the mature leaves of *Brachypodium*. We wish to thank Mike Jarvis for giving us the ^13^C NMR chemical shifts of his annealed flax and celery and for a draft version of his review.^48^ The UK High-Field Solid-State NMR Facility used in this research was funded by EPSRC and BBSRC (EP/T015063/1), as well as, for the 1 GHz instrument, EP/R029946/1. The work of RC, TK, RD and PD was supported by the UKRI grant underwriting the ERC advanced grant EVOCATE, Function and evolution of plant cell wall architecture for sustainable technologies EP/X027120/1. PKD was supported by EPSRC grant Bio-derived and Bio-inspired Advanced Materials for Sustainable Industries (VALUED), EP/W031019/1. AEP was supported by a Herchel Smith PhD scholarship by the University of Cambridge. The work of YY generating models of cellulose microfibril habits for conceptual analysis of glucose environments was supported as part of The Center for Lignocellulose Structure and Formation, an Energy Frontier Research Center funded by the US Department of Energy, Office of Science, Basic Energy Sciences, under Award number DE-SC0001090.

---

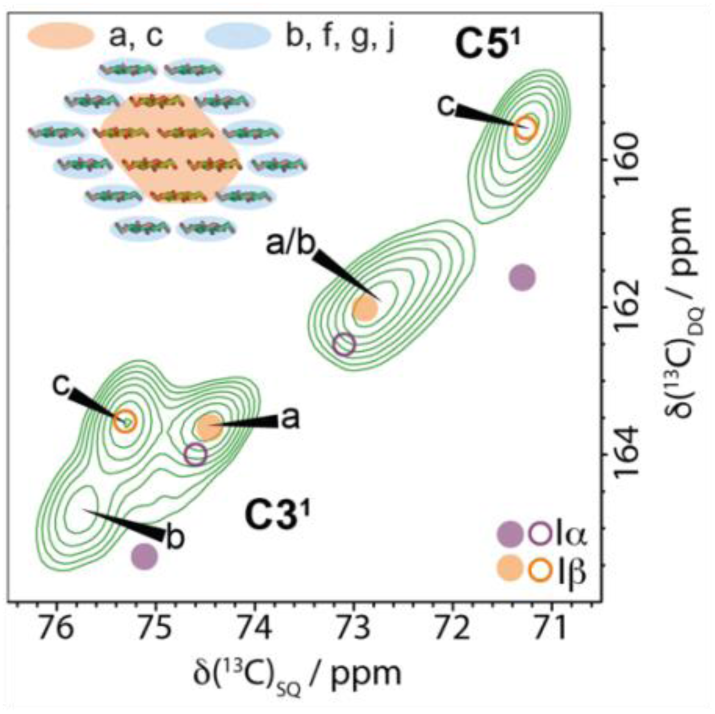
For Table of Contents only.

