## supplemental information file for "Using solid-state NMR to understand the structure of plant cellulose"

### Now at Department of Chemistry and Biochemistry, The Ohio State University, College of Arts and Sciences, 151 W. Woodruff Ave., Columbus, OH 43210, U.S.A.

#### Contents

#### Supplementary Figures

SI Figure 1: 30 ms $^{13}$C 2D CP PDSD spectrum of poplar wood.

SI Figure 2: TEM images of hCNFs of poplar wood.

SI Figure 3: Comparison of the 1D CP MAS NMR spectra of poplar wood, hCNFs of poplar wood and xylanase-treated hCNFs of poplar wood.

SI Figure 4: Cellulose of poplar wood maintains its structure in the fibrillation process to form hCNFs.

SI Figure 5: Cellulose fibril surface glucose environments in domain 2 are as well ordered as the glucose environments in domain 1 that include the fibril core.

SI Figure 6: The neutral carbohydrate region of the CP refocussed INADEQUATE showing the environments a and c from the xylanase treated hCNFs of poplar wood compared to the NMR shift positions of the two Iβ cellulose glucose units.

SI Figure 7: The C3-C1 and C3-C4 regions of a 30 ms $^{13}$C 2D CP PDSD spectrum showing the environments a and c from the xylanase treated hCNFs of poplar wood compared to the NMR shift positions of the two Iβ cellulose glucose units.

SI Figure 8: Neutral carbohydrate region of standard (blue) and water-edited CPMAS PDSD spectra (brown).

SI Figure 9: Similar water proximity of the three domain 2 surface glucose environments of cellulose.

SI Figure 10: Determining the interior to surface ratio from the relative amounts of glucose environment b and D2 versus core environments a and c.

**Supplementary Tables**

SI Table 1: The linewidths of glucose environments in cellulose from the C6 region of the CP INADEQUATE spectrum.

SI Table 2: Comparison of Iα and Iβ cellulose $^{13}C$ chemical shifts with assigned glucose environments a, b and c from spectral domain 1.

SI Table 3: Summary of solid-state NMR experimental parameters.

SI Table 4: Definitions of key terms used.

SI Table 5: List of acronyms

#### Supplementary Figures

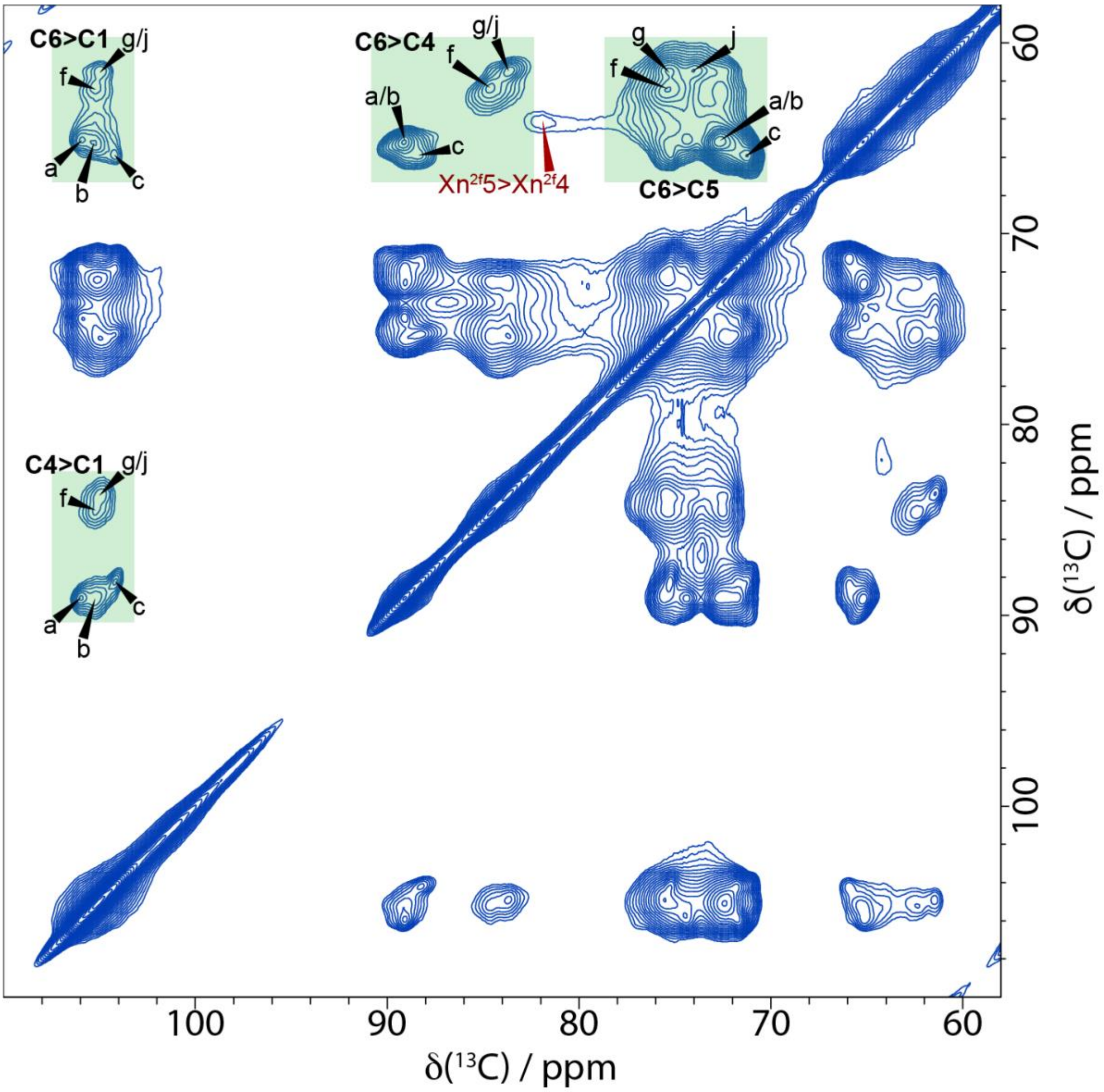


***SI Figure 1: 30 ms $^{13}$C 2D CP PDSD spectrum of poplar wood.*** *The main regions of the 30 ms $^{13}$C 2D CP PDSD spectrum where different glucose environments in cellulose can be resolved have been highlighted in green. The spectrum was recorded at a $^{13}$C Larmor frequency of 251.4 MHz and a MAS frequency of 12.5 kHz.*

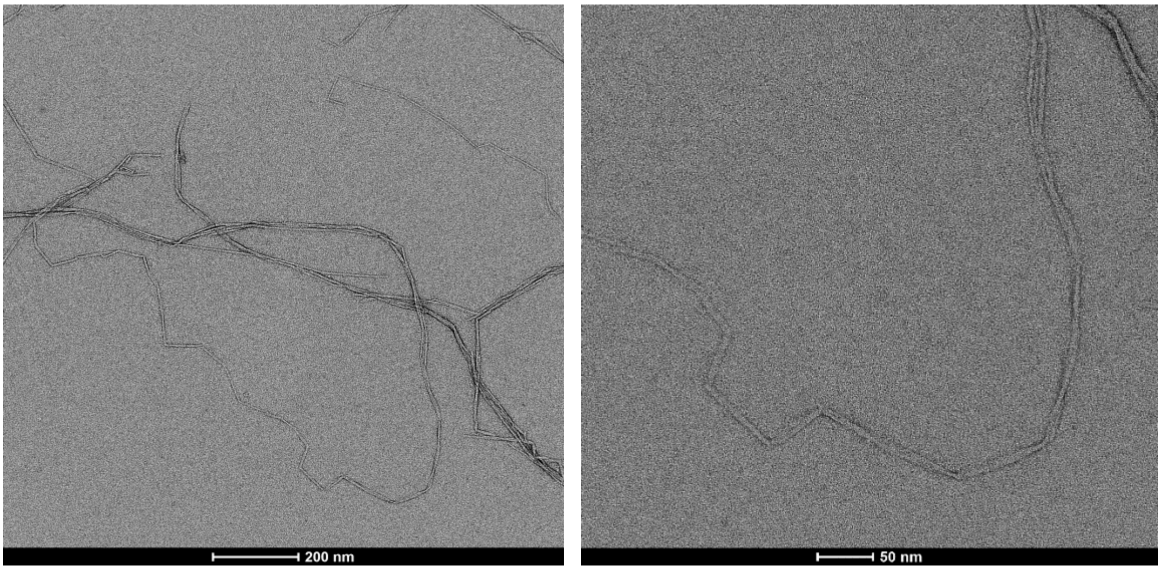

***SI Figure 2: TEM images of hCNFs of poplar wood.*** *The average width of microfibrils is ~3 nm.*

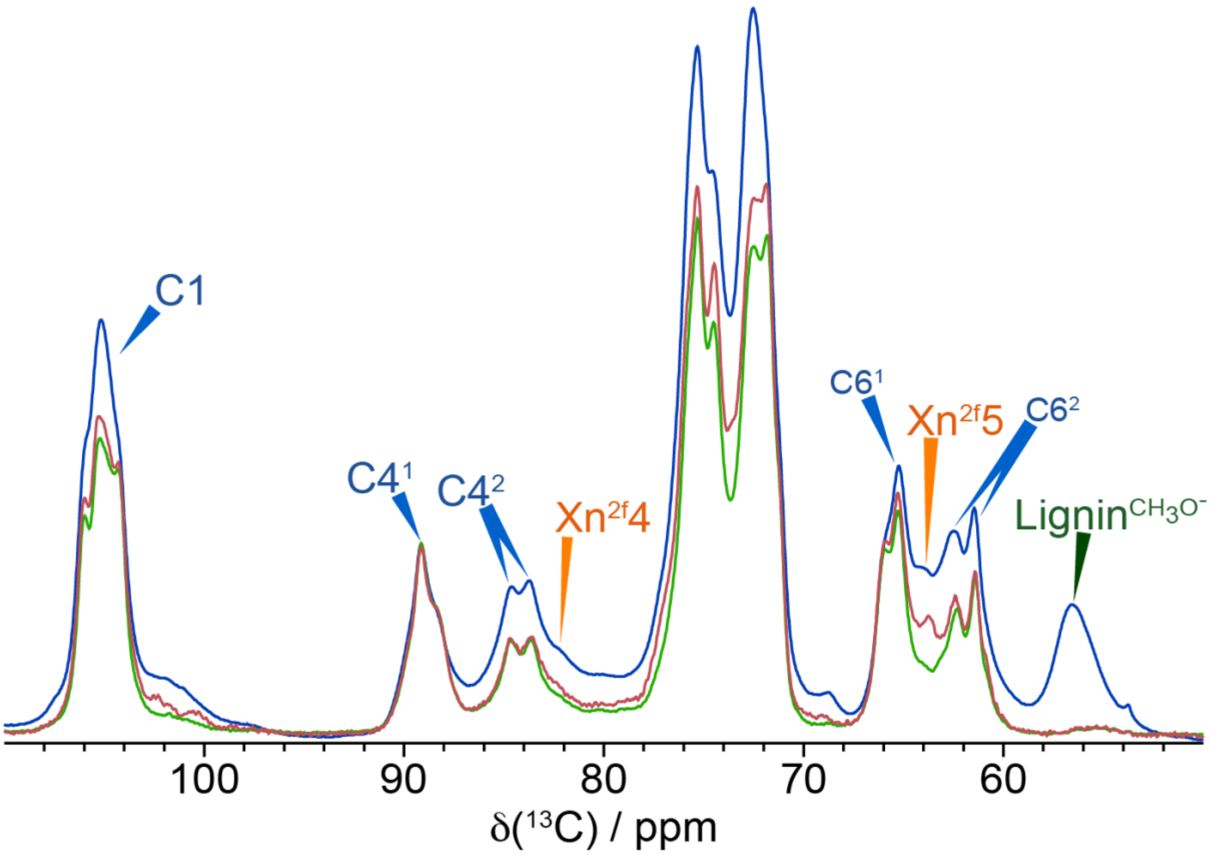


***SI Figure 3: Comparison of the 1D CP MAS NMR spectra of poplar wood, hCNFs of poplar wood and xylanase-treated hCNFs of poplar wood.*** *The comparison (normalised to C4 at 89 ppm) shows the difference in the 1D CP spectrum of poplar wood (blue), hCNFs of poplar wood when lignin and some hemicellulose has been removed (red) and xylanase-treated hCNFs of poplar wood where the majority of xylan has been removed (green). The production of hCNFs results in the removal of the broad background signal whilst the xylanase treatment removes more of the xylan. Spectra were recorded as in SI Figure 1.*

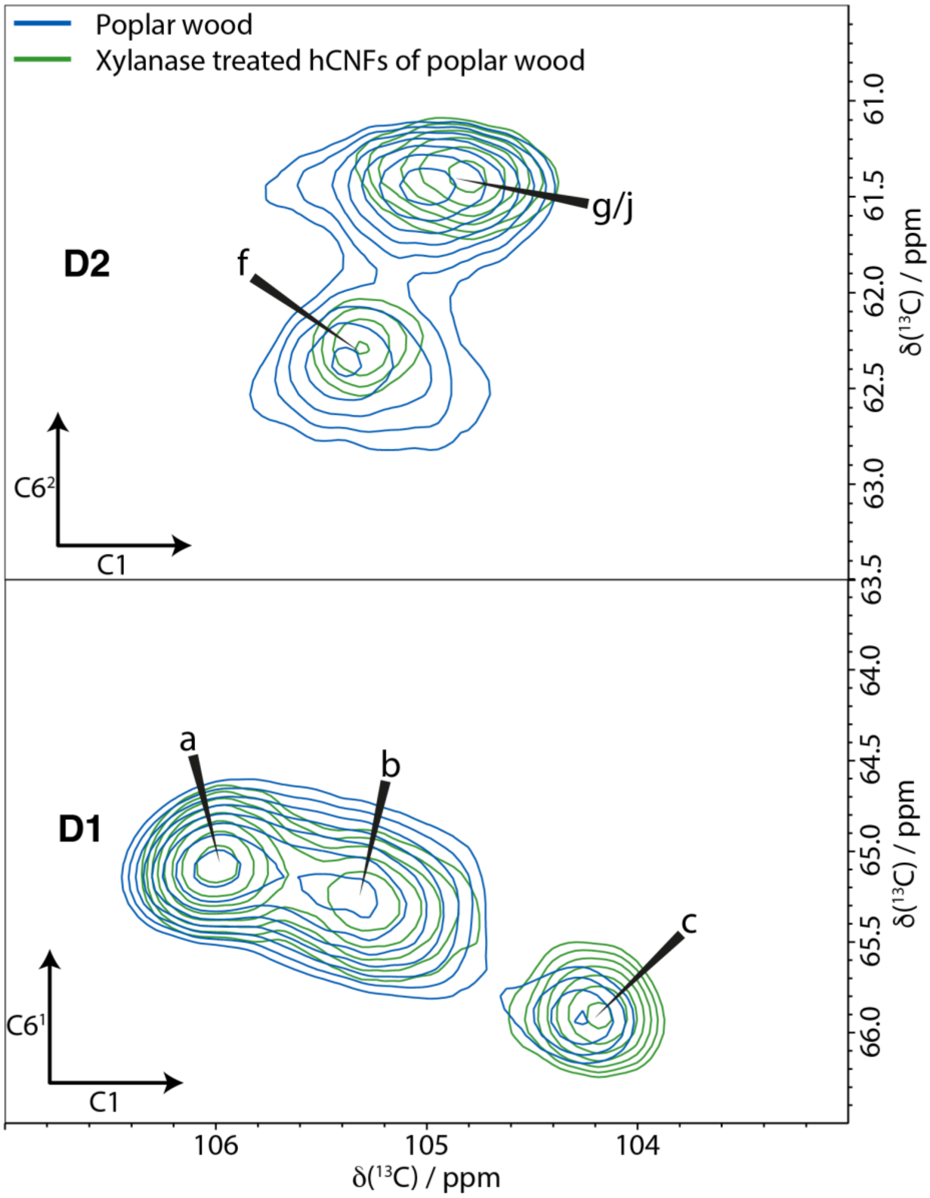


***SI Figure 4: Cellulose of poplar wood maintains its structure in the fibrillation process to form hCNFs.*** *Comparison of the C6-C1 region of 30 ms $^{13}C$ 2D CP PDSD spectrum of poplar wood (blue) and xylanase-treated hCNFs of poplar wood (green) normalised to the C61>C1 cross peak of site a . Spectra were recorded as in SI Figure 1.*

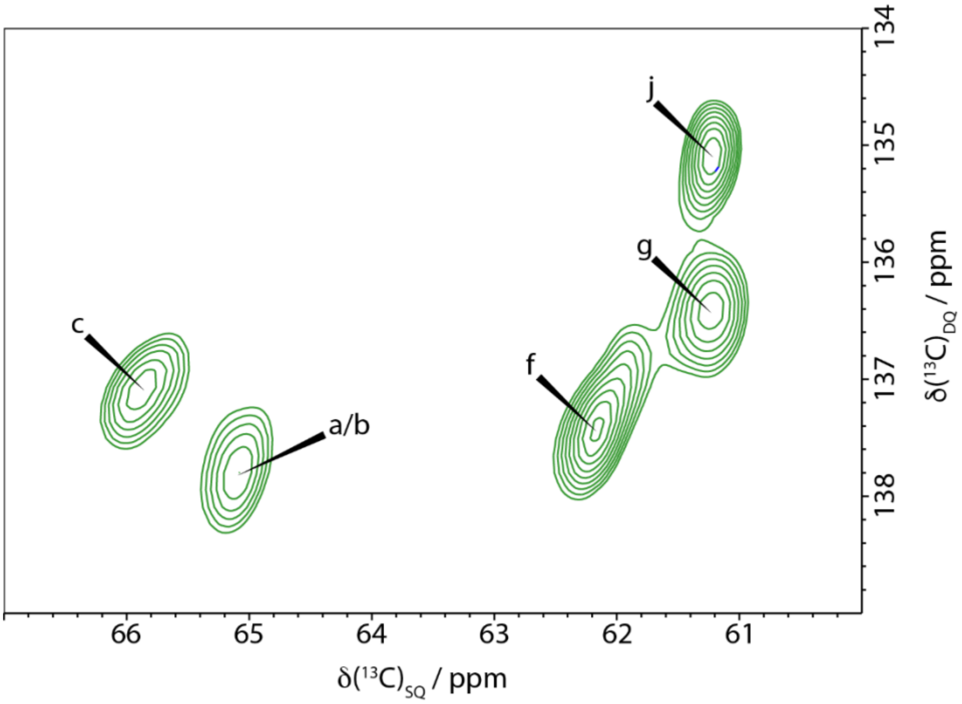


***SI Figure 5: Cellulose fibril surface glucose environments in domain 2 are as well ordered as the glucose environments in domain 1 that include the fibril core.*** *C6 region of the $^{13}C$ 2D CP refocused INADEQUATE MAS NMR spectrum of xylanase-treated hCNFs of poplar wood. The linewidths of the main domain 2 surface environments (f, g, j) are comparable to those of domain 1 (a, b, c) indicating significant (and similar) local order of the domain 2 glucose residues that lie on the fibril surface. Spectra were recorded as in SI Figure 1.*

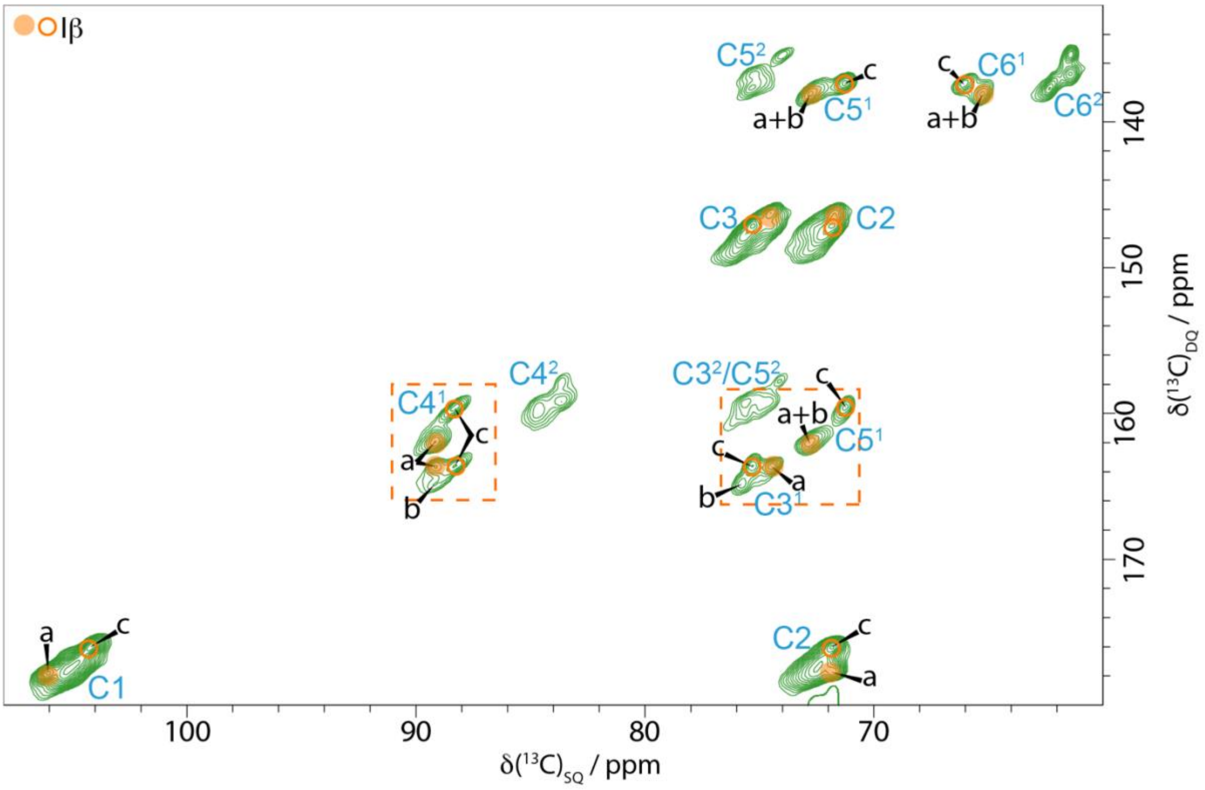


***SI Figure 6: The neutral carbohydrate region of the CP $^{13}$C refocussed INADEQUATE showing the environments a and c from the xylanase treated hCNFs of poplar wood compared to the NMR shift positions of the two Iβ cellulose glucose units.*** *Circles are the Kono et al. assignments[21,22] except for C1 which has been exchanged according to our assignments from the CP PDSD (see SI Fig. 7) with the chemical shifts corrected by Brouwer and Mikolajewski[23,57,58] This shows that cellulose sites a and c closely match all the corrected assignments for cellulose Iβ. The dashed boxes indicate the regions shown in figure 5. Spectra were recorded as in SI Figure 1.*

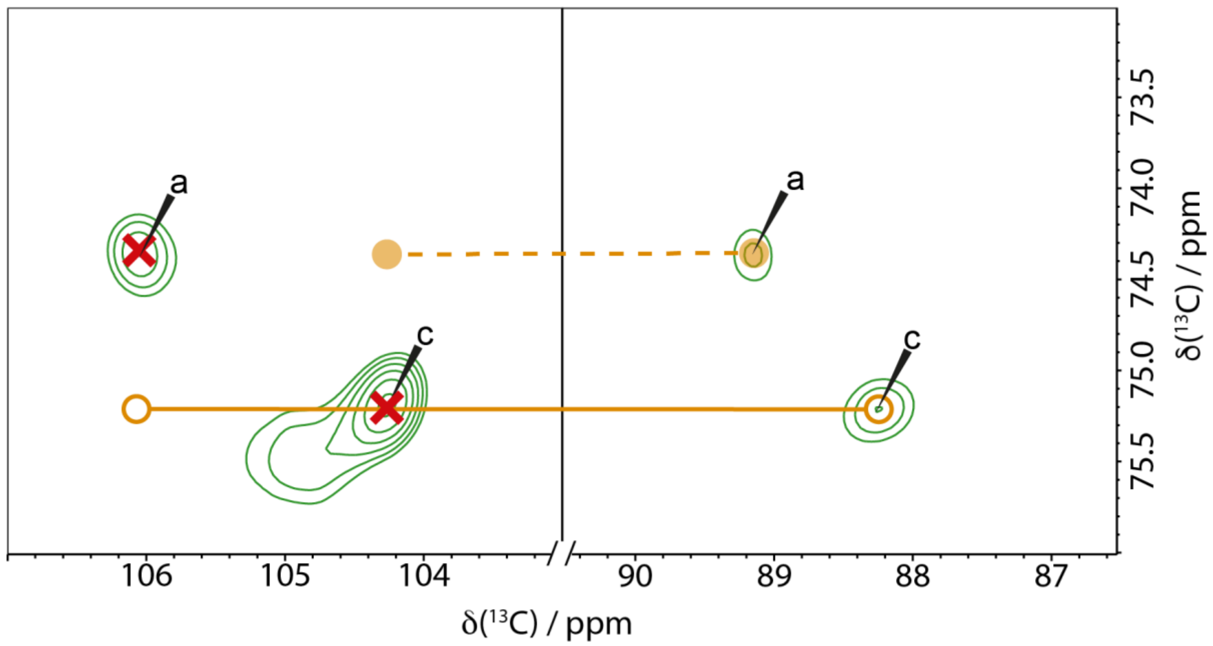


***SI Figure 7: The C3-C1 and C3-C4 regions of a 30 ms $^{13}C$ 2D CP PDSD spectrum showing the environments a and c from the xylanase treated hCNFs of poplar wood compared to the NMR shift positions of the two Iβ cellulose glucose units.*** *Circles are the Kono et al. assignments[21,22] with chemical shifts corrected by Brouwer and Mikolajewski.[23,57,58] The crosses indicate where the peaks would be if the C1 assignment were switched, as determined in this work, showing a and c closely match the corrected assignments for cellulose Iβ. Spectra were recorded as in SI Figure 1.*

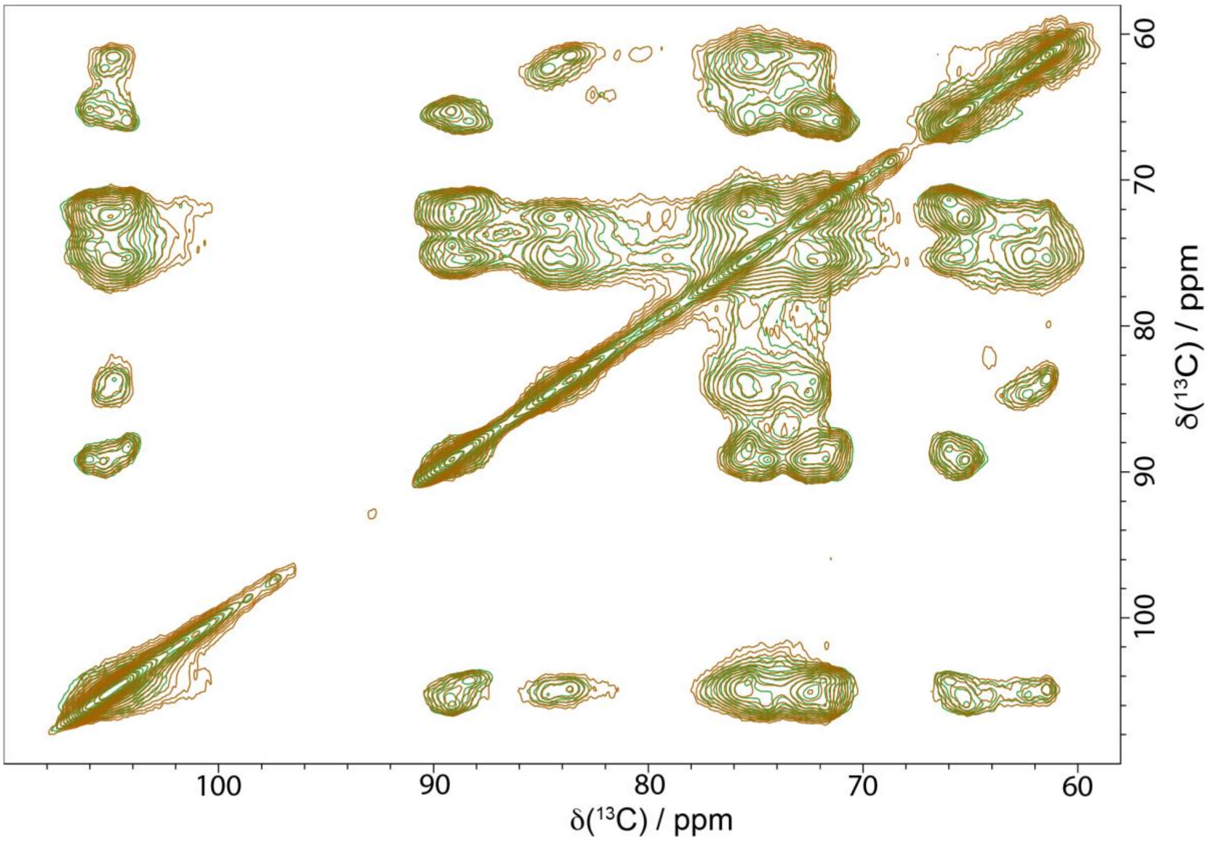


***SI Figure 8: Neutral carbohydrate region of standard (green) and water-edited $^{13}$C CPMAS PDSD spectra (brown).*** These spectra are normalised to the C1 diagonal peak at ~105 ppm. There is signal for all the cross peaks in both spectra however it is the relative intensities of these cross peaks that is most informative and is highlighted in Fig. 6.

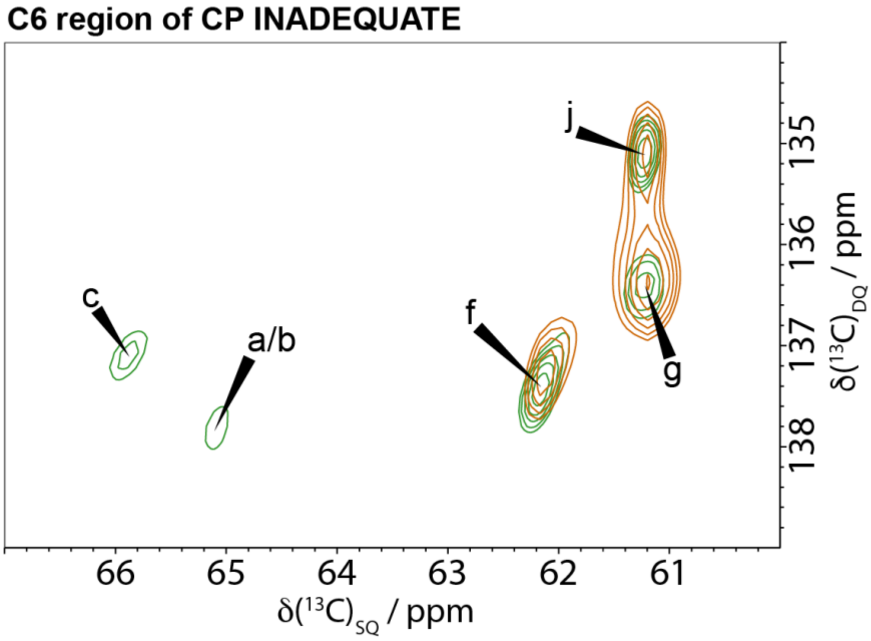


***SI Figure 9: Similar water proximity of the three domain 2 surface glucose environments of cellulose***. *A comparison of the C6 region of the standard (blue) and the water-edited (brown) CP refocussed INADEQUATE of the xylanase treated hCNFs of poplar wood and normalised to site a in the* C4[1]region of the CP refocussed INADEQUATE spectrum. *Spectra were recorded as in SI Figure 1*.

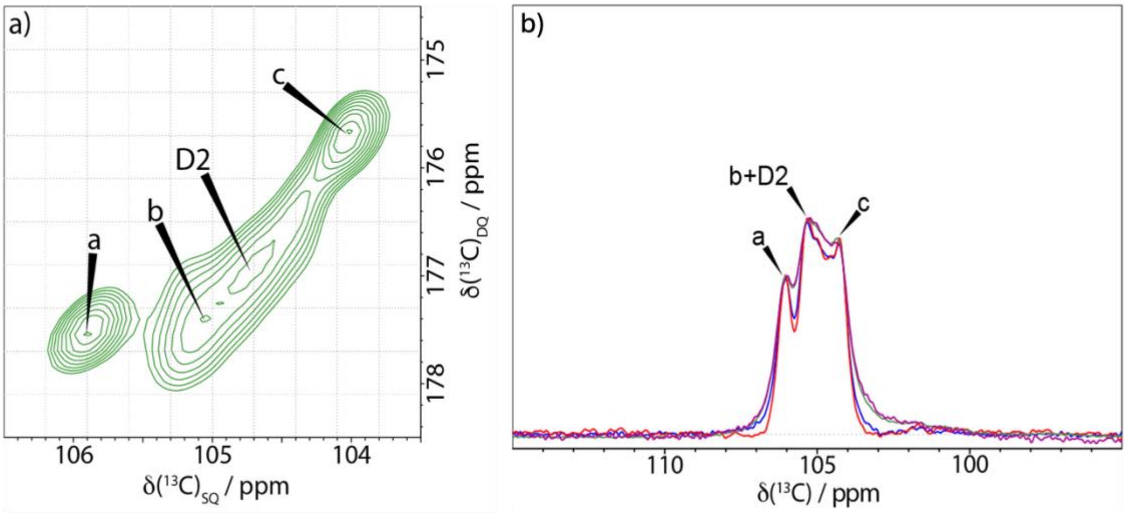


***SI Figure 10: Determining the interior to surface ratio from the relative amounts of glucose environment b and D2 versus core environments a and c.*** *a) C1 region of the $^{13}C$ 2D CP refocussed INADEQUATE MAS NMR spectrum of the xylanase treated hCNFs of poplar wood. This resolves fibril core sites a and c, allowing integration of the core environments separately from surface environments (b and domain 2). b) Comparison of the C1 region of the 1D CP (green), 1D quantitative DP (mauve) and two 1D CP Double Quantum filtered spectra with an echo time of 0.84 (red), or 2.24 ms (blue), of the xylanase-treated hCNF sample. Spectra were recorded as in SI Figure 1.*

#### Supplementary Tables

**SI Table 1: The linewidths of glucose environments in cellulose from the C6 region of the CP INADEQUATE spectrum of xylanase treated hCNFs of poplar stem.**

| Glucose environment | Linewidth C6 region CP INADEQUATE |
|---|---|
| a+b† | 0.57 ppm |
| c | 0.49 ppm |
| f | 0.57 ppm |
| g | 0.46 ppm |
| j | 0.41 ppm |

† The peak labelled a+b has a larger linewidth as it has two components that are ~0.2 ppm different in their NMR shift. Site f has a similar width to the peak for sites a + b and is elongated suggesting it could also be two environments which have yet to be resolved.

**SI Table 2: Comparison of Iα and Iβ cellulose $^{13}C$ chemical shifts with assigned glucose environments a, b and c from spectral domain 1.**

| | C1 | C2 | C3 | C4 | C5 | C6 |
|---|---|---|---|---|---|---|
| **Cellulose Iα** | | | | | | |
| Glucose Unit 1 (Down) † | 105.6 | 72.2 | 74.6 | 89.4 | 73.1 | 65.7 |
| Glucose Unit 2 (Up) † | 105.5 | 71.2 | 75.1 | 90.3 | 71.3 | 65.8 |
| **Cellulose Iβ** | | | | | | |
| Glucose Unit 1 (Centre) † | 106.1 | 71.7 | 75.3 | 88.4 | 71.4 | 66.0 |
| Glucose Unit 2 (Origin) † | 104.4 | 71.7 | 74.4 | 89.2 | 72.9 | 65.2 |
| **Cellulose Iβ (Corrected)** $ | | | | | | |
| Glucose Unit 1 (Centre) | 104.4 | 71.7 | 75.3 | 88.4 | 71.4 | 66.0 |
| Glucose Unit 2 (Origin) | 106.1 | 71.7 | 74.4 | 89.2 | 72.9 | 65.2 |
| **Native plant cellulose *** | | | | | | |
| a* (Cellulose Iβ origin) | 106.1 $ | 71.8 | 74.5 | 89.2 | 72.8 | 65.2 |
| b | 105.3 | 72.6 | 75.5 | 89.3 | 72.7 | 65.4 |
| c (Cellulose Iβ centre) | 104.3 | 71.8 | 75.4 | 88.4 | 71.3 | 66.0 |
| **Tunicate *** | | | | | | |
| a | 106.1 | 71.8 | 74.5 | 89.2 | 72.9 | 65.2 |
| c | 104.4 | 71.8 | 75.3 | 88.4 | 71.4 | 66.0 |

† Iα and Iβ cellulose $^{13}C$ chemical shifts as given by Brouwer and Mikolajewski[23]
$ Same shifts for cellulose 1β given by Brouwer and Mikolajewski[23] except the C1 have been switched based on our assignments.
* $^{13}C$ chemical shift assignments acquired in this work with an error of ± 0.1 ppm.

| SI Table 3 : Summary of Solid-state NMR experimental parameters | | | | | | | | |
|---|---|---|---|---|---|---|---|---|
| Sample | Experiment | Recycle Delay (s) | Echo/ Mixing time (ms) | Spectral Width (F2) (kHz) | Spectral Width (F1) (kHz) | Acquisition time (F2) (ms) | Acquisition time (F1) (ms) | Number co-added FIDs |
| Poplar wood | 1D CP | 2 | - | 100 | | 20 | - | 256 |
| | 1D DP | 2 | - | 100 | | 25 | - | 256 |
| | 1D DP | 20 | - | 100 | | 25 | - | 128 |
| | 2D CP refocussed INADEQUATE | 2 | 2.24 | 59 | 37.5 | 20 | 6.67 | 128 |
| | 2D CP PDSD | 2 | 30 | 75 | 37.5 | 25 | 8.2 | 64 |
| | 2D water edited CP refocussed INADEQUATE | 2 | $^{1}$H filter - 0.28<br>Diffusion time – 2<br>Mixing time – 2.24 | 28.5 | 100 | 20 | 4.7 | 270 |
| Xylanase treated hCNFs of poplar wood | 1D CP | 2 | - | 100 | | 20 | - | 256 |
| | 1D DP | 2 | - | 100 | | 25 | - | 256 |
| | 1D DP | 20 | - | 100 | | 25 | - | 128 |
| | 2D CP refocussed INADEQUATE | 2 | 2.24 | 59 | 37.5 | 25 | 6.7 | 128 |
| | 2D CP PDSD | 2 | 30 | 75 | 37.5 | 25 | 7.3 | 80 |
| | 2D CP PDSD | 2 | 200 | 75 | 37.5 | 25 | 5.1 | 80 |
| | 2D CP PDSD | 2 | 400 | 75 | 37.5 | 25 | 5.5 | 80 |
| | 2D Water edited CP PDSD | 2 | $^{1}$H filter - 0.32<br>Diffusion time – 2<br>Mixing time - 30 | 75 | 37.5 | 25 | 4.8 | 576 |
| | 2D water edited CP refocussed INADEQUATE | 2 | $^{1}$H filter - 0.28<br>Diffusion time – 2<br>Mixing time – 2.24 | 100 | 28.5 | 20 | 4.8 | 272 |

**SI Table 4: Definitions of key terms used.**

| Key terms | Definition |
|---|---|
| *tg*/Domain 1 spectral region | Refers to the glucose environments of cellulose with $^{13}C$ NMR chemical shifts of ~89 ppm for C4 and ~65 ppm for C6 ppm. |
| *gt* and *gg*/ Domain 2 spectral region | Refers to the glucose environments of cellulose with $^{13}C$ NMR chemical shifts of ~84 ppm for C4 and ~62 ppm for C6 ppm. |
| Hydroxmethyl (*tg*) conformation ratio | The ratio of the area under the spectrum of the two spectral domains in the C4 region at ~89 ppm and ~84 ppm |
| Glucose environment | A glucose residue in cellulose in a particular local environment/ position in the microfibril such that it has distinct NMR chemical shifts. |
| Amorphous cellulose (in NMR terms) | Cellulose that has no long-range order and is seen on NMR spectrum as a very broad component. |
| Crystalline cellulose (in NMR terms) | Cellulose with long range order (at least over a 5 Å length scale) and has very narrow line shape in the NMR spectrum. |
| Local order | NMR is sensitive to the local environment of a nucleus; local order refers to the short-range order that NMR probes (up to 0.5 nm). The line widths of the glucan environments in cellulose are similar suggesting similar short-range order for all the environments assigned in this work. Diffraction techniques require long-range periodic order to get high resolution patterns. |

**SI Table 5: List of Acronyms**

| Acronyms | Definition |
|---|---|
| NMR | Nuclear Magnetic Resonance |
| MAS | Magic Angle Spinning |
| CP | Cross Polarisation |
| DP | Direct Polarisation |
| PDSD | Proton Driven Spin Diffusion |
| INADEQUATE | Incredible Natural Abundance Double Quantum Experiment |
| hCNF | Holo-Cellulose Nanofibrils |
| D1 | Spectral Domain 1 |
| D2 | Spectral Domain 2 |
| TEM | Transmission Electron Microscopy |
